# Midkine Drives Reawakening of Dormant Melanoma Metastases

**DOI:** 10.1101/2025.05.27.656365

**Authors:** Carolina Rodriguez-Tirado, Thoufiqul Alam Riaz, Alcina A. Rodrigues, Nupura Kale, Ayax Perez-Gallegos, David Olmeda, Maria S. Soengas, Maria Soledad Sosa

## Abstract

A major challenge in melanoma treatment is the emergence of aggressive metastases from previously dormant disseminated cancer cells (DCCs). Here we identify Midkine (MDK) as a key factor that disrupts melanoma DCC dormancy by triggering an autocrine signal that suppresses the dormancy inducer NR2F1 and alters the p-p38/p-ERK ratio, via ALK signaling. Dormant melanoma DCCs in lymph nodes and visceral tissues exhibit an MDK^LOW^/NR2F1^HIGH^ phenotype. Transient NR2F1 activation for one month with a potent agonist counteracts MDK-driven mTORC1 activity and suppresses lung metastasis in mice bearing human melanoma DCCs. Dual targeting of MDK (genetic blockade) and NR2F1 (activation) markedly limits metastatic outgrowth and extends survival for 6 months in mouse models independent of BRAF status. These findings support the MDK-NR2F1 axis as a promising therapeutic target to sustain dormancy and prevent melanoma recurrence.

**Statement of Significance:** This work identifies Midkine as a key disruptor of tumor dormancy in melanoma, driving metastatic awakening via NR2F1 suppression. Dual targeting of Midkine (inhibition) and NR2F1 (activation) prolongs survival in preclinical models.

## INTRODUCTION

Disseminated cancer cells (DCCs) have long-been pursued as the source of metastatic relapses after the removal of primary tumors. A main feature of DCCs is that they can remain in a non-proliferative/quiescent state defined as dormancy until they reactivate and give rise to metastasis. Melanoma patients with detectable DCCs had an increased risk of dying of melanoma than patients without DCC detection^1,2^. In cutaneous melanoma, the timing for relapse could range from months to years^3^, yet the mechanisms that control dormancy and reactivation of melanoma DCCs are unclear. Here we explored a complex interplay between factors promoting metastasis and those maintaining dormancy in melanoma DCCs.

Recent publications on metastatic melanoma showed that the depletion of the secreted heparin-binding factor Midkine (MDK) reduced lymph node and lung metastases in mouse melanoma models. In addition, high levels of MDK correlated with a poor prognosis^4,5^. The described mechanisms of action for the pro-metastatic capacity of MDK to promote lymph node (LN) metastasis involved a paracrine signal from melanoma cells on lymphatic endothelial cells facilitating adhesion and transmigration between endothelial cells. Nevertheless, whether MDK could affect the metastatic outgrowth of disseminated melanoma cells in the lung in an autocrine manner remains unknown. The nuclear receptor NR2F1 is responsible for the dormancy phase of DCCs in head and neck, and breast cancers^6–8^. Moreover, NR2F1 expression in dormant DCCs from bone marrow aspirates from breast and prostate cancer patients is associated with their long-term dormancy^9,10^. Whether MDK modulates NR2F1 expression to promote reactivation of dormant DCCs has not yet been studied.

Here, we report that depletion of MDK in SK-Mel-147 melanoma cells (MDK^HIGH^), which carries NRAS Q61R mutation and is B-RAF wild-type, reduces lung metastatic burden resulting in a concomitant increasing in the frequency of dormant DCCs with high levels of NR2F1 expression. MDK, *via* an autocrine mechanism involving anaplastic lymphoma kinase (ALK) receptor signalling and inhibition of p38 (MAPK14) MAPK activity, suppresses NR2F1 expression and its target genes *SOX9*, *CDKN1B (P27KIP1)*, and *RARB*. Our pre-clinical results showed that treatment with an NR2F1 agonist reduced the size of lung metastatic foci formed by SK-Mel-147 MDK^HIGH^ cells by increasing the percentages of DCCs positive for NR2F1 and inhibiting mTOR signalling. Combining the NR2F1 agonist with MDK depletion further diminished lung metastasis size in mice injected with MDK^HIGH^ melanoma cells, regardless of BRAF mutation status (wild-type (WT) or V600E). This combination therapy significantly improved survival outcomes even if the treatment was given for only 1 month, providing a durable response of 5 months post-treatment in mice comparable to ∼10 years in humans. These findings highlight the therapeutic potential of dormancy-inducing strategies, such as NR2F1 activation alone or paired with MDK targeting, to reprogram melanoma cells and extend the dormant phase of MDK^HIGH^ melanoma DCCs and curb metastatic progression.

Lastly, supporting the physiological relevance of our findings, analysis of human lymph nodes and visceral tissues specimens reveals that dormant DCCs – negative for proliferation markers – exhibited significantly higher frequencies of the MDK^LOW^/NR2F1^HIGH^ putative dormant phenotype, with lower frequency of the putative reactivating MDK^HIGH^/NR2F1^LOW^ DCC phenotype.

Leveraging the discovery that MDK silences NR2F1 to drive metastatic reawakening, we offer a promising pre-clinically actionable dual-target therapy that forces melanoma cells via a long-lasting reprogramming, into prolonged dormancy, eradicating their metastasis-initiating capacity. Notably, this approach appears effective in the BRAF-mutant context as well as the NRAS-mutant setting, for which specific NRAS-directed inhibitors are not yet established in clinical practice.

## RESULTS

### MDK depletion induces NR2F1 expression in melanoma lung DCCs while preventing metastasis formation

We have previously reported that in melanoma patients, high levels of the growth factor MDK was associated with worse overall and disease-free survival^4,5^. Mice subcutaneously (s.c.) injected with melanoma cells with high levels of MDK (MDK^HIGH^) formed lung metastases whereas mice injected with MDK^LOW^ melanoma cell lines did not^5^. Moreover, downregulation of MDK in MDK^HIGH^ melanoma cells showed reduced spontaneous lung metastases. We and others have previously shown that the nuclear receptor NR2F1 is responsible for the dormancy phase of, among other cancer types, head and neck squamous carcinoma cells^6^. Further, high NR2F1 levels in minimal residual disease stage have been linked to better survival in prostate and breast cancer patients^6,9,10^. Thus, we hypothesized that MDK^LOW^ melanoma cell lines, which do not form lung metastasis^5^, might have high levels of NR2F1. To this end, we measured the levels of NR2F1 by immunofluorescence (IF) in disseminated cancer cells (DCCs) in lungs of mice s.c. injected with MDK^HIGH^ (SK-Mel-147, SK-Mel-28, and 451LU) and MDK^LOW^ (WM35 and WM164) melanoma cell lines. Of note, SK-Mel-147 cells expressed the highest protein levels of MDK when compared with the other five cell lines^5^. We observed that DCCs from the MDK^HIGH^ SK-Mel-147 melanoma cells had ∼20% DCCs positive for NR2F1 (**Fig. 1a&b)** and SK-Mel-28, and 451LU cells (**Supp. Fig. 1a**) had ∼45%. By contrast, DCCs from the MDK^LOW^ melanoma cell lines WM164 (**Fig. 1a)** and WM35 (**Supp. Fig. 1a**) were ∼65% positive for NR2F1. Interestingly, the MDK^LOW^ WM35 cell line, like the WM164 cells, is poorly metastatic when compared to MDK^HIGH^ cell lines^5^. Therefore, high MDK levels correlate with higher metastatic potential and with lower NR2F1 levels in DCCs.

**Figure 1.**
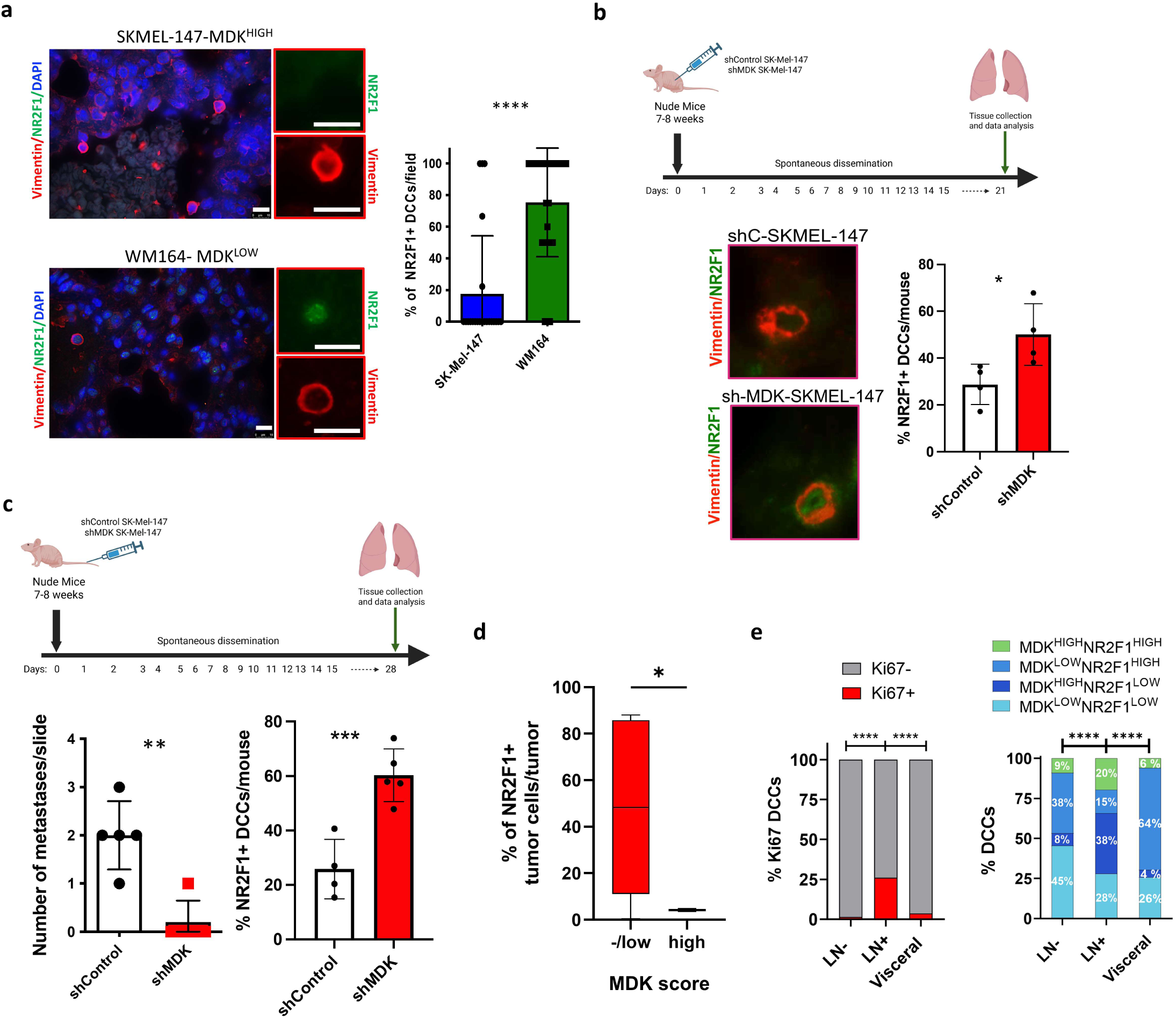
Midkine represses NR2F1 levels in melanoma DCCs. **a.** SK-Mel-147 (SK) and WM164 (WM) cells were s.c. injected into nude mice. Lung sections were stained for NR2F1 (green), human Vimentin (red, recognizes only human vimentin) and DAPI (blue). Zoom in pictures show single DCCs. 10 μm scale bars (white bars). Bar plots show percentage of NR2F1+ DCCs/field. Mann-Whitney test. **b.** GFP-SK-Mel-147 (shControl and shMDK) cells were implanted s.c. in nude mice. Lungs were collected 21 days later and embedded in paraffin. Lung sections were stained for NR2F1 (green), Vimentin (red) and DAPI (Blue). Percentage of NR2F1+ DCCs/mouse in lungs injected with shControl- or shMDK-SK-Mel-147 (two shRNAs against MDK) cells. Mann-Whitney test. **c.** shControl and shMDK GFP-SK-Mel-147 cells were i.v. injected in nude mice. Lungs were collected and embedded in paraffin. Number of metastasis was determined on H&E stained sections and the percentage of NR2F1+ DCCs/mouse was determined after IF. Mann-Whitney test. **d.** Primary tumors were scored for MDK based on IHC staining (**Suppl.** Fig. 1). Sequential sections for each tumor were stained for NR2F1 and HMB45. The percentage of NR2F1+ tumor cells per tumor sample (N=18) was plotted. **e.** Lymph nodes carrying metastasis (LN+, N=8 patients, number of DCCs analysed: 122), LN without apparent metastasis (LN-, N=7 patients, number of DCCs analysed: 77) and visceral metastatic tissues (lung, brain, and omentum, N=4 patients, number of DCCs analysed: 248) were used to identify percentages of positive or negative Ki-67 DCCs (left panel) and the percentages of DCCs with the indicated phenotypes among patients (right panel). Fisher’s exact test (P<0.0001).

Previous work showed that MDK depletion in SK-Mel-147 cells reduced the frequency of nude mice developing lung metastases compared to controls after s.c. injections^5^. To investigate whether depletion of MDK could affect spontaneous dissemination to the lungs, we examined lungs at an earlier time point - 21 days after s.c. injection - when macrometastases have not yet formed, and quantified the overall DTC burden. Strikingly, the total lung DCC load (detected by GFP and human vimentin staining, with the latter not cross-reacting with mouse vimentin) was comparable between mice injected with SK-Mel-147 shControl and shMDK cells (**Fig. 1b, Suppl. Fig. 1a–d**), indicating that MDK knockdown did not impair dissemination to the lungs. Consistent with this, we obtained similar results in nude mice injected with MDK^HIGH^ SK-Mel-28 cells (**Suppl. Fig. 1e**).

To isolate the direct effect of MDK on metastatic growth without influence from the primary tumor, we performed experimental metastasis assays. We observed less metastatic load in mice injected with GFP-SK-Mel-147 shMDK cells when compared to shControl cells (**Fig. 1c**). This result suggests that loss of MDK impairs reactivation of dormant DCCs, limiting their metastatic outgrowth. Interestingly, we observed an increased percentage of NR2F1 positive DCCs in mice s.c. (∼50% average) or i.v. (∼60% average) injected with SK-Mel-147 shMDK cells when compared to the SK-Mel-147 shControl group (∼25% average) (**Fig. 1b&c, Suppl. Fig. 1b&c**). Similar results were obtained when MDK^HIGH^ SK-Mel-28 cells were depleted of MDK (**Suppl. Fig. 1f**). We conclude that the loss of MDK prevents lung metastasis formation via a putative mechanism dependent on the induction of NR2F1 in DCCs. As predicted, NR2F1 levels (mRNA and protein) were increased upon MDK depletion in cultured melanoma cells (**Suppl. Fig. 1g &h**) arguing that active MDK signaling is needed to repress NR2F1 expression. Interestingly, growing lung metastases from spontaneous and experimental metastasis assays were negative/low for NR2F1 when compared to DCCs (**Suppl. Fig. 1b&c**), which is consistent with previous studies and confirms the association of NR2F1 expression with cell cycle arrest^6–8^. Importantly, the NR2F1 levels in non-metastatic WM164 cells with low levels of MDK expression^5^ compared to metastatic MDK^HIGH^ SK-Mel-147 cells were downregulated upon addition of recombinant human MDK (rhMDK) (**Supp. Fig. 1i**). These results suggest that MDK signaling controls NR2F1 expression in melanoma cells.

### Human melanoma DCCs can display a MDK^LOW^/NR2F1^HIGH^ dormancy-like phenotype

Next, we examined the expression levels of NR2F1 and MDK in human samples from primary melanomas. We determined MDK levels in primary melanoma by IHC as shown before^5^ and classified the samples into positive or negative scores (**Supp. Fig. 1j)**. Next, in a consecutive section, we stained with NR2F1 and HMB45 (melanoma specific marker) antibodies (**Supp. Fig. 1k**). We observed that MDK^LOW/NEGATIVE^ melanomas were in average 45% (range from 15-80%) NR2F1 positive, whereas MDK^HIGH^ melanomas were only ∼5% positive for NR2F1 (**Fig. 1d**).

We next examined the co-expression of MDK and NR2F1 in individual DCCs detected in lymph node (LN) and visceral biopsies from melanoma patients. LN samples were divided into two groups based on histopathology: those with DCCs but no detectable macro-metastasis (LN-) and those containing macro-metastases (LN+). Visceral lesion biopsies were derived from the peripheral margins of metastases from lung, brain, and omentum. Among DCCs, Ki-67 expression was found in 25% of LN+ cells, compared with only 1.4% in LN- and 3% in visceral lesions samples (**Fig. 1e**). These results suggest that LN- for macrometastasis and visceral DCCs are more quiescent than DCCs in LN+ for metastasis, even though in the latter group they were still found as single DCCs.

We then assessed MDK and NR2F1 expression patterns within the same biopsies and identified four distinct DCC phenotypes: MDK^LOW^/NR2F1^HIGH^ (putative dormant), MDK^HIGH^/NR2F1^LOW^ (putative reactivating), MDK^HIGH^/NR2F1^HIGH^ (double high) and MDK^LOW^/NR2F1^LOW^ (double low). The putative dormant phenotype comprised 38% of LN- DCCs and 64% of visceral DCCs, but only 15% of LN+ DCCs (**Fig. 1e & Supp. Fig. 1l&m**). In contrast, the putative reactivating MDK^HIGH^/NR2F1^LOW^ DCC phenotype was most enriched in LN+ samples (38%) and markedly lower in LN- (8%) and visceral (4%) DCCs (**Fig. 1e & Supp. Fig. 1l&m**).

When focusing specifically on Ki-67- DCCs, the dormant and double low phenotypes represented the majority of DCCs found (>80%), while reactivating and double high phenotypes accounted for less than 10% (**Supp. Fig. 1n**). These observations suggest a strong association between DCC with lack of or reduced proliferation and the putative dormant MDK^LOW^/NR2F1^HIGH^ state, with marginal contribution from putative reactivating MDK^HIGH^/NR2F1^LOW^ phenotype. The MDK^LOW^/NR2F1^LOW^ population represents half of the total of LN- DCCs and 26% in visceral DCCs, which were largely Ki-67- (**Fig. 1e**, ∼97%), suggesting that the lack of MDK expression could support a dormant/low proliferation phenotype that is independent of NR2F1. Notably, MDK^LOW^/NR2F1^LOW^ population was reduced by half in LN+ samples compared to LN- (**Fig. 1e**, right panel). Conversely, the presence of MDK^HIGH^/NR2F1^HIGH^ (double high) cells suggests that MDK may not always repress NR2F1, or that these cells represent a transient state transitioning from the MDK^LOW^/NR2F1^HIGH^ dormant phenotype toward the reactivating MDK^HIGH^/NR2F1^LOW^ state. Further studies are required to elucidate the biological relevance of these double high and double low subpopulations.

Overall, these results suggest the existence of MDK^LOW^/NR2F1^HIGH^ (putative dormant) and MDK^HIGH^/NR2F1^LOW^ (putative reactivating) phenotypes in human DCCs from melanoma biopsies, which may predict the progression of the melanoma disease.

### MDK depletion induces cell cycle arrest by upregulating NR2F1 via ALK receptor and p38 signalling

A feature of cellular dormancy is the presence of high p38 (SAPK) activity and low ERK (MAPK) activity^11^. Thus, we calculated the phospho-p38(p-p38) to phospho-ERK (p-ERK) ratio in shControl and shMDK SK-Mel-147 cells. We showed a higher p-p38/p-ERK ratio in SK-Mel-147 shMDK when compared to the SK-Mel-147 shControl cells (**Fig. 2a**). These results suggest that downregulation of MDK may induce a quiescence phenotype as reported^11^ . To determine whether the upregulation of NR2F1 upon MDK depletion was dependent on p38 activity, we knocked down total p38 protein by using siRNA in shMDK SK-Mel-147 cells. We observed that p38 depletion downregulated NR2F1 protein levels (**Fig. 2b**). Moreover, treatment of shControl SK-Mel-147 cells with SB203580 (a p38α/β inhibitor) and TAK371 (p38α inhibitor) also decreased NR2F1 levels and p-MK2, a surrogate for p38 activity (**Fig. 2c**). The levels of SOX9, RARB and CDKN1B, downstream targets of NR2F1, were increased in SK-Mel-147 shMDK cells when compared to shControl cells (**Fig. 2a&d**). Moreover, an MDK inhibitor (iMDK) increased *NR2F1*, *RARB* and *CDKN1B* levels in SK-Mel-147 cells **(Supp. Fig. 2a**). Overall, these results suggest that MDK depletion in SK-Mel-147 MDK^HIGH^ melanoma cells upregulates NR2F1 levels and its targets downstream of p38 signalling.

**Figure 2.**
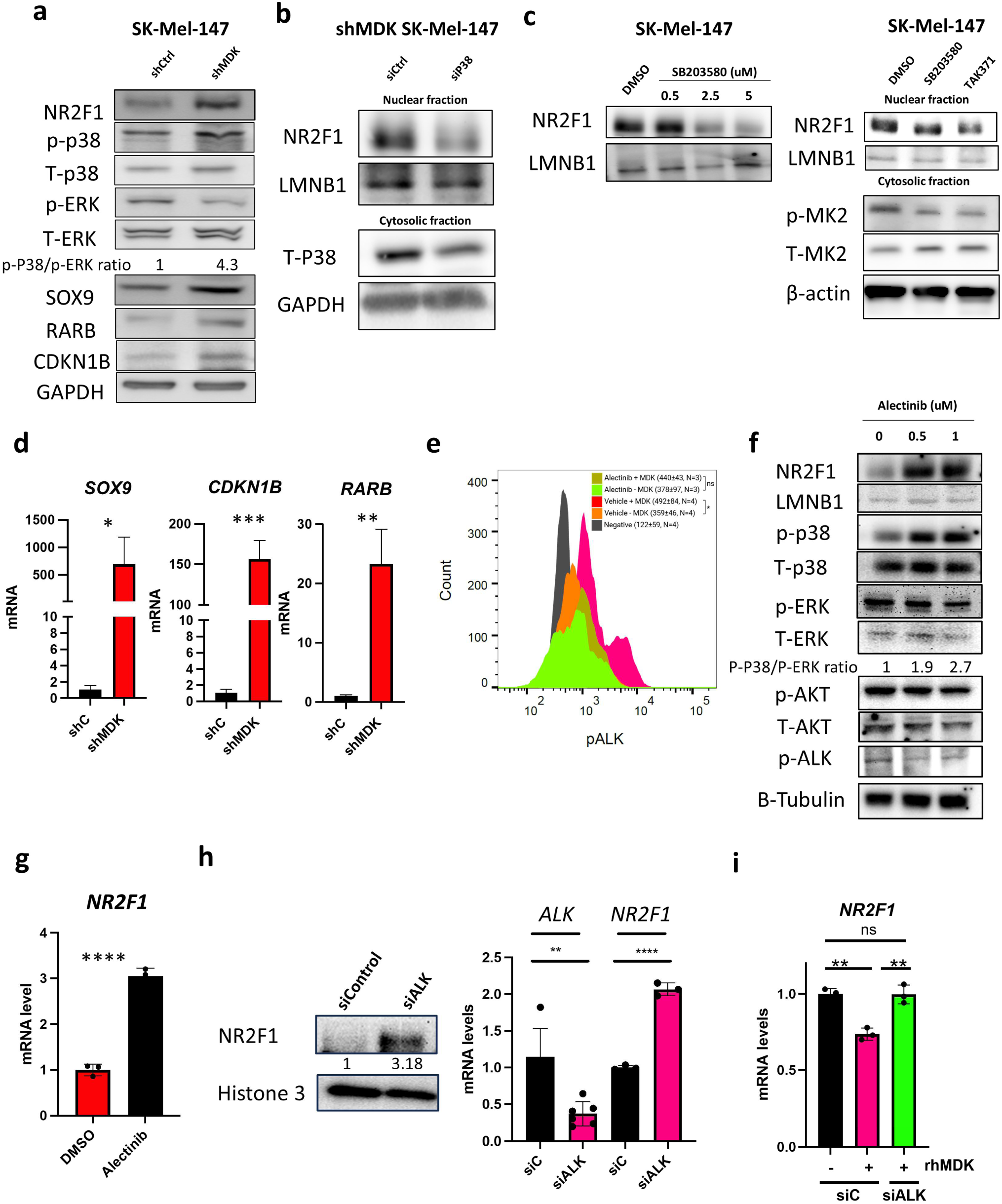
Midkine represses NR2F1 and its downstream targets via p38 and ALK. **a.** Western blot for NR2F1, p-p38, p38, p-ERK, ERK, SOX9, RARβ, CDKN1B and GAPDH from shControl and shMDK SK-Mel-147 cells. The ratio of p-p38 to p-ERK was calculated by densitometry. **b.** Western blot for NR2F1 in the nuclear fraction and total p38 in the cytosolic fraction from shMDK SK-Mel-147 cells transfected with siC/sip38 (75nM, 48h). Lamin B1 and GAPDH are loading control proteins. **c.** SK-Mel-147 cells were seeded in 2D and then treated with SB203580 (dose curve, left panel) and with 5 µM and 0.5 µM TAK715 (right panel) for 24 hours in serum-free media. Western blots for the indicated targets are shown. MK2 (phosphorylated and total) levels were measured as a p38 target. **d.** SOX9, RARB, and CDKN1B mRNA levels were measured by qRT-PCR from shControl and shMDK SK-Mel-147 cells. Unpaired t-test**. e.** Intracellular flow cytometry for phosphoALK in SK-Mel-147 cells treated with Alectinib and rhMDK. Mean+SD from 3-4 independent experiments are shown (One-Way ANOVA). **f&g.** SK-Mel-147 cells were seeded in 2D and then treated with Alectinib for 24 hours. western blots (**f**) and qRT-PCR (**g**) for the indicated targets are shown. Nuclear extraction was used to detect NR2F1. The ratio of p-p38 to p-ERK was calculated. Unpaired t-test**. h.** SK-Mel-147 cells were seeded in 2D and then transfected with siControl and siALK for 48 hours. qRT-PCR and western blots for the indicated targets are shown. Nuclear extraction was used to detect NR2F1. Numbers represent densitometry analysis for the NR2F1 band over Histone 3. Unpaired t-test**. i.** siControl and siALK SK-Mel 147 cells were daily treated with rhMDK or vehicle for 48 h and qRT-PCR levels for *NR2F1* were measured. Unpaired t-test.

To determine whether MDK had similar effects in a previously validated dormancy model, we used the dormant head and neck squamous cell carcinoma (HNSCC) D-HEp3 cells^6^. We previously published that dormant D-HEp3 cells have high levels of NR2F1 when compared to the proliferative T-HEp3 cells, from which they were derived^6^. We measured MDK levels by IF and observed that proliferative T-HEp3 cells have higher percentage of MDK+ cells than D-HEp3 cells (**Supp. Fig. 2b**). Thus, the inverse correlation between MDK and NR2F1 levels also exists in this HNSCC model, with the MDK^LOW^/NR2F1^HIGH^ profile associated with a dormancy phenotype. Treatment of D-HEp3 cells with rhMDK decreased NR2F1 levels (**Supp. Fig. 2c**). Similarly, treatment of SK-Mel-147 cells with rhMDK decreased NR2F1 levels (**Supp. Fig. 2c**). Next, we decided to overexpress MDK in D-HEp3 cells and determine their ability to exit dormancy using the chorionallantoic membrane (CAM) system as described^6^ (**Supp. Fig. 2d**). We observed that exogenous expression of MDK allowed dormant D-HEp3 cells to exit dormancy and proliferate (**Supp. Fig. 2d**).

We next decided to target ALK, a known receptor for MDK, using the ALK inhibitor alectinib^12–14^. We validated that SK-Mel-147 cells express ALK by flow cytometry at comparable levels with NB-EBC1, a previously described human neuroblastoma cell line expressing WT ALK (**Supp. Fig 2e**). Addition of rhMDK to SK-Mel-147 cells increased phosphorylation of ALK **(Fig. 2e)**. Notably, alectinib was able to block rMDK-dependent phosphorylation of ALK in SK-Mel-147 cells without changes in total ALK (**Fig. 2e** and **Suppl. Fig. 2f**). This effect was also observed in response to zotizalkib, a more potent ALK inhibitor and without changes in total ALK (**Supp. Fig 2g**). Next, we observed that alectinib-treated SK-Mel-147 cells upregulated nuclear NR2F1 protein and mRNA levels and increased phosphorylation of p38 while downregulating phosphorylation of ERK and ALK when compared to control-treated cells (**Fig. 2f-g** and **Supp. Fig. 2h**). Similar results were obtained in murine B16R2L cells, which express high levels of MDK^4^ (**Supp. Fig. 2i**). We then silenced human ALK using siRNA, efficiently reducing mRNA abundance, cell surface receptor abundance, and phosphorylation of the receptor after rhMDK stimulation (**Fig. 2h** and **Sup. Fig. 2j**). After confirming efficient ALK knockdown, we observed an increase of NR2F1 mRNA levels and nuclear protein levels (**Fig. 2h**). The expression levels of NR2F1 remained unchanged when additional MDK receptors^14^, specifically low-density lipoprotein receptor-related protein (LRP1), neurogenic locus notch homolog protein 2 (NOTCH2), syndecan-3 (SDC3), and Protein Tyrosine Phosphatase Receptor Zeta 1 (PTPRZ1) were silenced (**Sup.** **Fig**. **2k**). Importantly, the addition of rhMDK to SK-Mel 147 cells repressed NR2F1 (as shown in **Supp. Fig. 2c**) but this effect was blocked in the absence of ALK expression (**Fig. 2i** and **Supp. Fig. 2l**), suggesting that MDK represses NR2F1 expression via ALK signaling.

We next measured whether MDK inhibition induces cell cycle arrest. To this end, we transduced SK-Mel-147 cells with a cyclin dependent kinase 2 (CDK2) biosensor (CDK2 DHB-mVenus). The cytosolic to nuclear ratio of the fluorescent reporter informs whether cells are in G0/G1 (nuclear only), S (both) or G2/M phases (cytosolic only)^15^. We observed an increase (2-fold change) in the percentage of cell arrested in G0/G1 phase upon MDK depletion (**Fig. 3a**). Concomitantly, the percentage of cells in S+G2/M phases was decreased upon MDK depletion (**Supp. Fig. 3a**). We also performed cell cycle analysis by Ethynyl-2’-deoxyuridine (EdU) incorporation and flow cytometry and showed that depletion of MDK in SK-Mel-147 cells increased the percentage of cell arrested in G0/G1 phase from ∼40% (in control group) to ∼80% (**Fig. 3b** and **Supp. Fig. 3b**).

**Figure 3.**
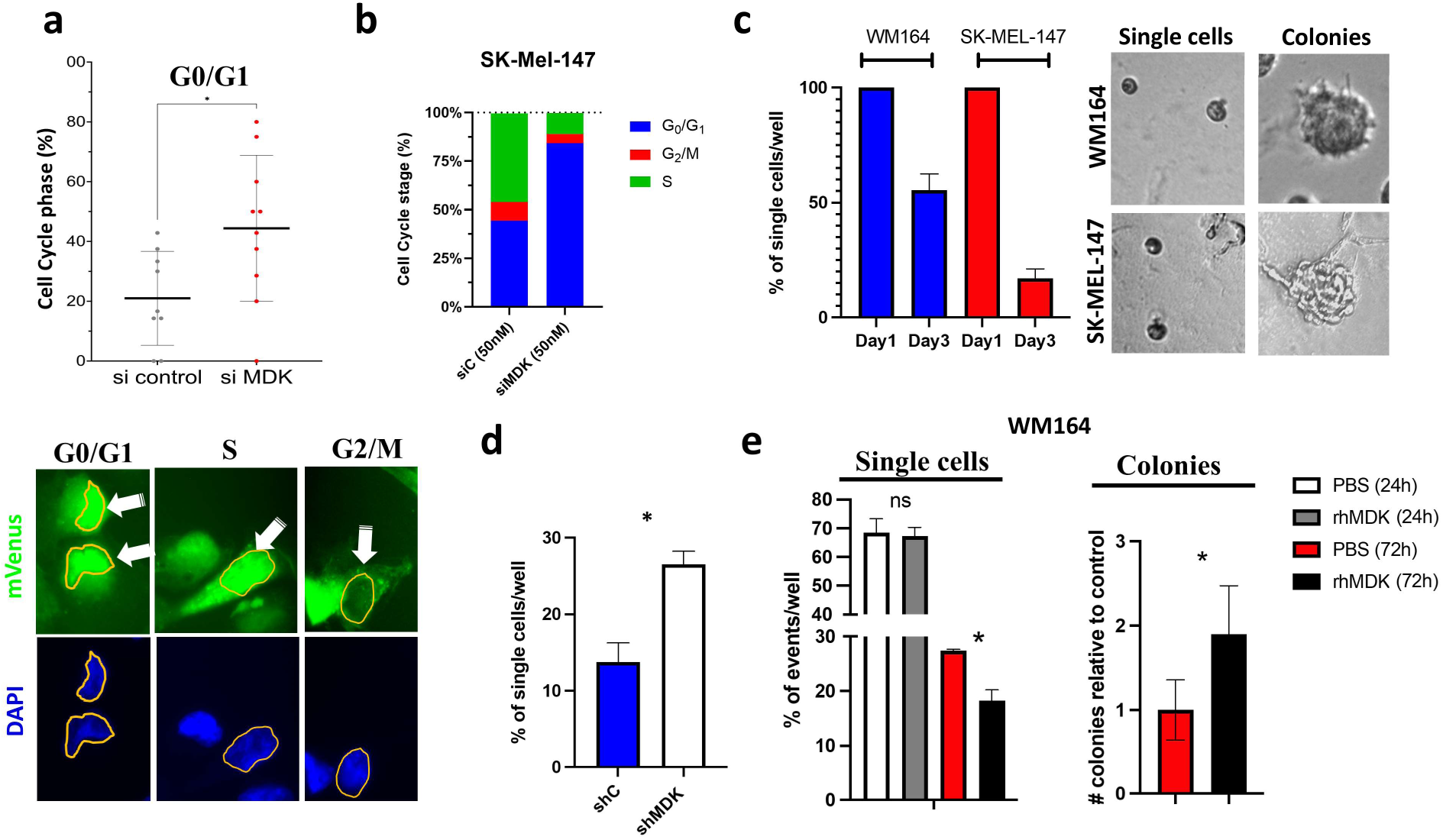
Midkine depletion induces cell cycle arrest. **a.** SK-Mel-147 cells stably expressing mVenus reporter were transfected with siControl or siMDK and after 48 hours cells were fixed and the percentage of cells in G0/G1 phases was plotted. Representative images for the indicated cell cycle phases are shown. Unpaired t-test. **b.** SK-Mel-147 cells were transfected with siMDK or siControl. Forty-eight hours later cell cycle analysis was done by FACS. **c.** SK-Mel-147 and WM164 cells were seeded on 3D Matrigel. Number of single cells per wells at day 1 and 3 was counted. Representative images for single cells and colonies are shown. **d.** Percentage of single cells on day 3 in shControl vs. shMDK SK-Mel-147 cell lines. Mann-Whitney. **e.** WM164 cells were seeded in 3D Matrigel, 24 h later cells were treated with PBS (white bars) and rhMDK (grey bars, 20 ng/ml) for 24h and with PBS (red bars) and rhMDK (black bars, 20 ng/ml) for 72 h. Percentages of single cells and colonies were graphed. 3 wells/condition, number of events counted=50-100. Representative of two experiments. Unpaired t-test.

Next, we compared the behaviour of MDK^HIGH^ vs. MDK^LOW^ cells in 3D Matrigel cultures by seeding them at single-cell density to model solitary DCC behaviour. Over time, the fraction of single cells, which is initially 100%, declines as these cells begin to proliferate and form new colonies. We observed that, after 3 days of culture, the percentage of remaining single cells was ∼50% for WM164 (MDK^LOW^) and ∼15% for SK-Mel-147 (MDK^HIGH^) cells, arguing that WM164 cells have lower proliferation rate when compared to SK-Mel-147 cells (**Fig. 3c**). We then depleted MDK in SK-Mel-147 cells and observed that, 3 days after seeding, a higher percentage of melanoma cells remained as single cells (∼25%) in shMDK group when compared to shControl cells (∼15%) (**Fig. 3d**). Similarly, we observed that WM164 cells treated with rhMDK, which reduced NR2F1 levels (**Supp. Fig. 1i**), resulted in a reduction of the percentage of single cells and an increase in colonies formed after 4 days, arguing for an increase in proliferation (**Fig. 3e**). We conclude that MDK depletion upregulates an NR2F1-dependent dormancy program via ALK and it mediates cell cycle arrest.

### Agonist-driven NR2F1 activation prevents melanoma metastasis

Next, we decided to evaluate the effect of an NR2F1 agonist (compound 26, C26), previously described by us^7^, on dormancy-regulating genes in MDK^HIGH^ cells. This agonist binds the ligand binding domain of NR2F1, activates its transcriptional activity which upregulates the NR2F1 gene itself and triggers transcription of downstream targets for dormancy induction^7^. Treatment of MDK^HIGH^ SK-Mel-147 cells with C26 showed upregulation of NR2F1, RARB, and CDKN1B (**Fig. 4a**) whereas it did not change MDK secreted protein levels (**Supp. Fig. 4a**). Similar results were obtained in two additional MDK^HIGH^ melanoma cell lines, SK-Mel-103 and SK-Mel-28 cells (**Supp. Fig. 4b&c**).

**Figure 4.**
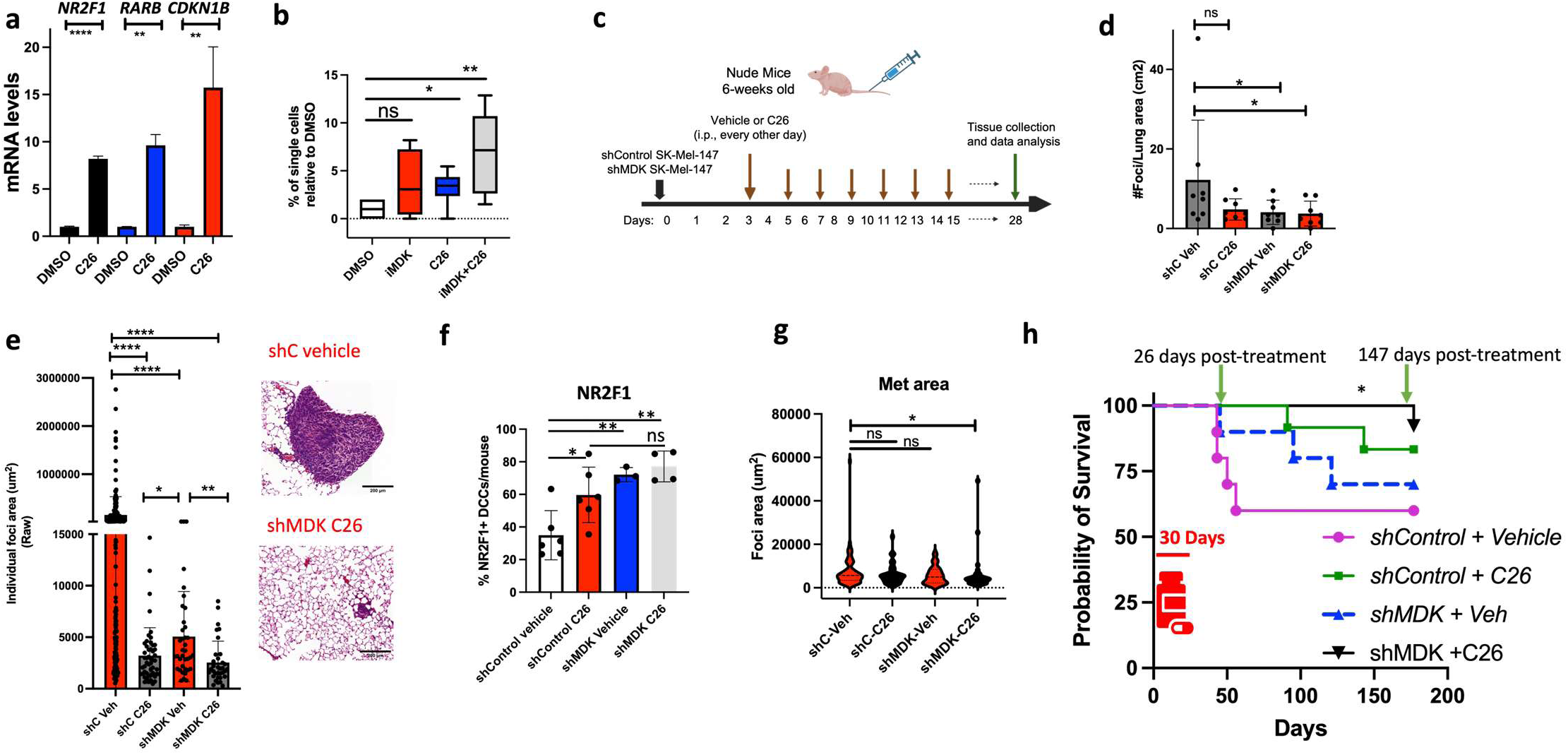
**Dual targeting of MDK (inhibition) and NR2F1 (activation) blocks metastasis and prolongs survival in preclinical models**. **a.** SK-Mel-147 cells were treated with 1.0 µM C26 for 48h. Relative mRNA levels of NR2F1, RARB and CDK1NB was plotted. Unpaired t-test. **b**. SK-Mel-147 cells were seeded in Matrigel and treated with vehicle, compound 26 and MDK inhibitor. Quantification of single cells at 5 days after respective treatment groups. N= 4 experiments. Mann-Whitney test. **c.** Schematic model for the i.v. injection of shControl and shMDK SK-Mel-147 cells into nude mice followed by treatment with vehicle or C26 for one month. **d&e.** Collected lung as shown in **c** were stained by H&E and number (**d**) and size (**e**) of metastasis were measured. Representative images of macrometastasis and micrometastasis are shown. Scale bar= 200 um. **f.** Percentage of lung NR2F1+ DCCs in each condition. N= ∼200 DCCs/group, 4-6 animals/group. **g**. shControl and shMDK SK-Mel-28 cells were i.v. injected (one million cells/mouse) into nude mice followed by treatment with vehicle or C26 for one month. Collected lung were stained by H&E and size of metastasis were measured. N=5 mice/group. Mann Whitney. **h.** Same schematic model as in **c** was followed for the treatment. After one month treatment, mice were euthanized when morbidity signs were observed. N=8 mice/group. Gehan-Breslow-Wilcoxon test.

We then evaluated in MDK-expressing melanoma cells changes in cell cycle-related genes, known to be regulated by NR2F1^6,7^, in response to C26 treatment. We observed that while rhMDK reduced the expression of CDKN1B, CDKN1A, CDKN2B, CDKN2A and CDKN2C, the treatment with C26 induced these genes in SK-Mel-147 cells (**Supp. Fig. 4d&e**). Expression of the CDKN2D gene did not respond to rhMDK nor C26 in SK-Mel 147 cells (**Supp. Fig. 4d&e)**. Similar results were obtained in SK-Mel-28 cells treated with C26, except that we observed no effect on CDKN1B gene, however there was upregulation of CDKN2D (**Supp. Fig. 4f**). Thus, loss of MDK or activation and induction of NR2F1 is associated with key CDK inhibitor upregulation.

To determine whether these changes translated to C26 inducing cell cycle arrest, we treated 3D cultures of MDK^HIGH^ SK-Mel-147 cells with C26 alone or in combination with the iMDK. We observed that when used individually, the iMDK- and C26-treated groups had ∼3-fold increase in the percentage of single cells when compared to control group, arguing that the treatments kept cells arrested. Interestingly, the combination of both drugs increased 7-fold the percentage of single cells when compared to control condition (**Fig. 4b**). This result led us to evaluate in a pre-clinal model whether the combination of MDK inactivity together with NR2F1 activation could serve as a potential therapeutic strategy to prevent metastasis. To this end, nude mice intravenously injected with shControl or shMDK GFP-SK-Mel-147 cells were randomly divided into vehicle or C26 treatment groups 3 days post-injection and treated for one month (**Fig. 4c**). We concluded that the number of metastatic foci in the lungs was reduced upon MDK depletion consistent with our previous results^5^ and that the combination of MDK depletion plus C26 also significantly decreased the number of metastasis when compared to control group (**Fig. 4d**). The number of metastases was also reduced by treating animals injected with shControl cells with C26 alone, but this was not statistically significant. However, when we looked at the size of the individual foci, we observed that C26 alone potently reduced the foci size when compared to shControl vehicle-treated group. This was also true for inhibition of MDK plus C26, which further reduced the size of metastatic foci when compared to MDK depletion effect alone (**Fig. 4e**). Interestingly, analysis of the percentage of NR2F1+ DCCs, 30 days post-treatments, showed an increased in the percentage of NR2F1+ DCCs in all conditions when compared to control group (**Fig. 4f**). There were no differences in the percentages of NR2F1+ DCCs between the three treatments, however, a trend towards higher percentages of NR2F1+ DCCs was observed in MDK depletion plus C26 group (**Fig. 4f**).

Next, we decided to evaluate the in vivo effect of C26 on metastasis formation in a BRAF mutant context. We first confirmed that depletion of MDK in SK-Mel28 cells (V600E BRAF) increased NR2F1 levels (**Supp. Fig. 4g**) and that stimulation by exogenous rhMDK decreased nuclear NR2F1 intensity when compared to control group by IF (**Supp. Fig. 4h**). Next, we i.v. injected SK-Mel 28 cells into nude mice and followed the same experimental design as in **Fig. 4c**. We observed that although there was a trend towards a decrease in number of metastases in C26-, shMDK-, C26+shMDK-groups when compared to control group, the statistical analysis was not significant (**Supp. Fig. 4i**). Similarly, the size of metastases was not significantly different in C26- and shMDK-groups when compared to control group, however, we observed a decrease in sizes when mice were treated with C26 and injected with shMDK-SK-Mel-28 cells (**Fig. 4g**). These results suggest that combination of MDK inactivation with NR2F1 activation may be a potential therapeutic strategy to reduce metastatic burden in BRAF mutant melanomas.

To evaluate how the effects above translated to a survival benefit, we treated animals for only the first month as in **Fig. 4c**, and then measured survival rates over time. We observed that mice injected with shMDK SK-Mel-147 cells had better survival rates (90%) than control group (∼60%) 26 days post-treatment (**Fig. 4h**). However, this difference disappeared at 147 days post-treatment (60% shControl+vehicle vs. 70% shMDK+vehicle) (**Fig. 4h**). Importantly, the group injected with shMDK SK-Mel-147 cells plus C26 had higher survival rates (∼90%) than shControl + vehicle and shMDK + vehicle groups at 147 days post-treatment (**Fig. 4h**). Mice injected with shControl SK-Mel-147 cells plus C26 had a trend towards better survival rates (∼80%), however, this was not statistically significant when compared to shControl + vehicle group. These results, support that dual targeting of MDK by inhibition and NR2F1 via agonists could have a significant impact (5 months post-treatment in mice which equals to ∼10 years in humans) on preventing metastatic disease even after cessation of the treatment.

### Agonist-driven NR2F1 activation blocks mTOR-dependent MDK signaling and prevents melanoma metastasis

To understand how the NR2F1 agonist alone or combination with MDK depletion could inhibit metastatic growth, we evaluated mTOR activity, previously shown to be activated by MDK in a paracrine manner on lymphatic endothelial cells^5^. In addition, we had published evidence that treatment of HNSCC with C26 in *in vitro* settings inhibited the mTOR pathway^7^. Consistent with this, we observed reduced phospho-S6 (pS6) levels in SK-Mel-147 cells treated with C26 compared to control cells in vitro (**Fig. 5a**). Overexpression of NR2F1 in SK-Mel-147 cells also decreased pS6 (**Suppl. Fig. 5a**). We also observed increase phosphorylation of AMPKα at threonine 172 upon C26 treatment, which was inhibited in NR2F1 depleted cells (**Suppl. Fig. 5b&c**), indicative of a metabolic response triggered by NR2F1 activation and associated with inhibition of mTOR signalling. Interestingly, the levels of pS6 in single DCCs from the lungs isolated from shControl vehicle vs. shControl C26 groups (from **Fig. 4c**) showed no significant differences (**Supp. Fig. 5d**). We hypothesized that this result might reflect the biology of dispersed single DCCs in the lung epithelium, which may not be actively proliferating. However, this could change in areas where multiple DCCs are found in close proximity (high density DCC and cluster areas). To test this hypothesis, we examined lungs from the shControl vehicle group and found that only 20% of isolated DCCs were positive for pS6 (**Fig.5b**). In contrast, in areas with clusters and high density of DCCs of more than 7 DCCs, the proportion of pS6-positive cells increased significantly to 70% (**Fig. 5b**). This suggests that clustered DCCs may be exiting dormancy and engaging in proliferation. Importantly, analysis of LN biopsies revealed that DCC clusters exhibited higher pS6 signal compared to solitary DCCs (**Fig. 5c**). These results validate in human samples our mouse findings and suggest that pS6 activation may be necessary for the transition to a proliferative state. Importantly, MDK depletion or C26 treatment reduced the percentage of pS6+ DCC clusters compared to the vehicle control (**Fig. 5d & Suppl. Fig. 5e**).

**Figure 5.**
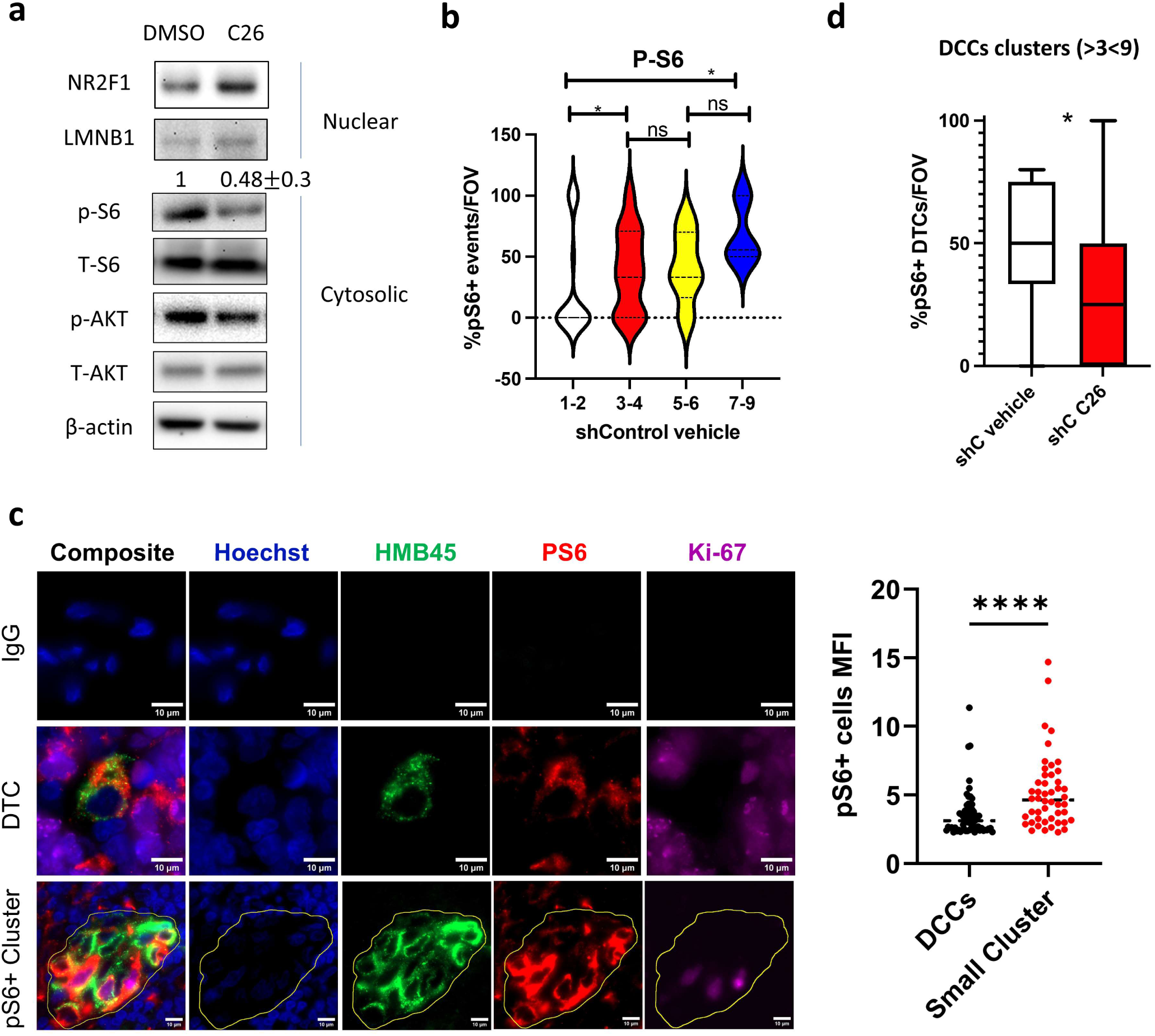
Agonist-driven NR2F1 activation blocks mTOR-dependent MDK signaling. **a.** SK-Mel-147 cells were treated with C26 for 24 h and nuclear and cytosolic fractions were collected and plotted against the indicated antigens. **b.** Collected lung from shControl +vehicle group as shown in Fig. 4c were stained by IF for vimentin and pS6. The percentage of DCC positive for pS6 is shown in each group. N= 6 mice. **c.** pS6 signal in solitary DCCs (N= 66) vs. DCC clusters (3-9 cells, N=48 total cells) from LN+ biopsies was plotted. N= 18 patients. Representative pictures of solitary DCC and clusters of DCCs. **d**. DCC cluster with more than 3 DCC and less than 9 DCCs per field of view (FOV) were stained for pS6 and the percentage of positive cells was plotted for the indicated groups. N= 6 mice/group.

## DISCUSSION

Previous studies have shown the role of MDK in favouring cancer growth, survival, angiogenesis, and the formation of distant metastasis^4,5^. However, the mechanism of action of MDK when DCCs are lodged at secondary organs has not been studied previously. Here, we reveal an NR2F1-driven dormancy program in melanoma N-ras and BRAF mutant DCCs that is repressed by the growth factor MDK in an autocrine manner.

Previous studies demonstrated that reducing MDK levels in the SK-Mel-147 cell line led to a decrease in lymphangiogenesis both at the primary tumor site and in LN. Interestingly, this MDK reduction did not impact the tumor onset and the formation of blood vessels in primary tumors^5^. Our findings reveal that depleting MDK in primary tumors does not impair dissemination of DCCs to the lungs. This observation may suggest that in this model, the spread of cancer cells to distant organs still can occur through blood vessels when lymphangiogenesis is dysfunctional. This is consistent with clinical data showing that completion lymph node dissection after a positive sentinel node does not improve melanoma-specific or overall survival, and instead primarily enhances regional disease control, arguing for new strategies to contain melanoma DCC at distant sites^16^.

MDK is a secreted cytokine that travel to distant sites, perhaps in exosomes, from the primary tumor^5^. In this study, we have used experimental metastasis by tail vein injection to eliminate the effect of a pre-metastatic niche created by the primary tumor. Nevertheless, we still observed the lack of metastatic capacity of DCCs depleted of MDK. This result suggests that the autocrine MDK activity could be enough to trigger metastatic outgrowth of DCCs, as depletion of MDK in DCCs impaired metastasis formation.

Our previous research has generated promising results in preventing metastasis formation using the NR2F1 agonist C26 in a pre-clinical model of HNSCC^7^. This success lends support to the concept that therapies aimed at inducing dormancy could be advantageous in certain types of cancer. A particularly noteworthy finding emerged in this work when we administered C26 to mice harboring MDK^HIGH^ DCCs with mutant-Ras and wild-type BRAF background. While the C26 treatment alone did not significantly alter the number of metastatic events per lung, it resulted in a marked reduction in the size of metastatic lesions (**Fig. 4**) arguing it contained awakening. This suggests that activating NR2F1 with C26 may hinder the MDK-driven growth program during the early phases of DCC reawakening (from single DCC to clusters of DCCs). This observation aligns with our findings in human LN DCCs (**Fig. 5**), which indicate activation of the mTOR pathway during the transition from single DCC to small clusters of DCCs. Importantly, C26 appears to reduce the mTOR activation in clusters of DCCs. These results are particularly significant as they pave the way for investigating the potential use of mTOR inhibitors, either alone or in combination with C26, to prevent the reactivation of MDK^HIGH^ DCCs in melanoma patients. Noteworthy, a new clinical trial with patients diagnosed with breast cancer patients showed that combined treatment inhibiting autophagy and mTOR targeted dormant DCCs and prevent late recurrence (CLEVER pilot trial, NCT03032406).^17^

Our study revealed that approximately 80% of animals injected with MDK^HIGH^ shControl-SK-Mel-147 cells and treated with C26 alone for one month remained disease-free after 5 months post-treatment, compared to the control group. However, this difference was not statistically significant likely due to the number of animals.

Notably, only the combination of MDK depletion and C26 treatment significantly improved survival rates for 5 months past a one-month treatment. These result support a long-lasting response to C26 treatment, potentially reflecting epigenetic reprogramming, consistent with our observation that NR2F1 regulates histone mark landscapes in dormant cancer cells^6^. This data may become critical for translational research in where the design of a treatment for melanoma patients carrying high levels of MDK will imply to not only stop the pro-tumorigenic signal driven by MDK but also enforce a long-lasting dormancy state via NR2F1 activation. Previous studies have shown the development of nanoparticles containing mRNA against MDK that successfully work in pre-clinical models of hepatocellular carcinomas^18^. Therefore, a combination of anti-MDK nanoparticles and the NR2F1 agonist could potentially serve as a therapeutic strategy to prevent metastasis in MDK^HIGH^ melanomas.

In vivo experiments using MDK^HIGH^/BRAF^V600E^ SK-Mel-28 cells showed that only the combination of MDK depletion and C26 treatment significantly reduced metastasis size compared to the control. Interestingly, C26 alone did not statistically decrease the number or size of metastases in these cells. This is different from the results in N-ras Q61R BRAF wild-type SK-Mel-147 cells, where C26 alone decreased the size of metastases. Perhaps, other signals may be blocking NR2F1 expression in a BRAF mutant context.

In patients diagnosed with melanoma, the presence of a limited number of DCCs in LN- has been reported, and their presence is associated with increased risk for death when compared to patients without DCC detection or low density of DCCs^1,2^. Importantly, the removal of sentinel lymph nodes does not extend overall survival of patients with melanoma, suggesting that DCCs in other organs may play important roles in survival rates. Using the same HMB45 antibody as previously described to identify single DCCs in LNs^1,2^, we conclude that dormant DCCs (negative for proliferation marker) found in LN- and distant organs biopsies predominantly present the putative dormant MDK^LOW^/NR2F1^HIGH^ phenotype while the putative reactivating MDK^HIGH^/NR2F1^LOW^ phenotype is scarce.

From a translational perspective, these findings could be validated in LN biopsies collected at the time of melanoma diagnosis with follow up data on date of recurrence and/or death. Interestingly, our exploratory and underpowered analysis of a small cohort of patients (N=10) with follow up information (**Suppl. Table 1**) shows a trend, where 3 out of 4 patients that were deceased in less than 3 years carried a dominant putative reactivating MDK^HIGH^/NR2F1^LOW^ DCC population. On the other hand, 6 out of 6 patients that remained alive 6-19 years later or recurred after 7 years carried a putative dominant dormant MDK^LOW^/NR2F1^HIGH^ DCC population and/or dominant double negative population. These results are consistent with our hypothesis but remain preliminary and should be interpreted as exploratory. We propose that the ratio between MDK and NR2F1 may serve as a biomarker for predicting lethal recurrence in melanoma patients.

In recent years, the role of MDK has been implicated in the modulation of both innate and adaptive immune cells, primarily demonstrated using murine B16-derived cells in immunocompetent animals^4^. While our current study using athymic mice may limit our ability to investigate immune system interactions, our previous work has shown that NR2F1 can induce dormancy similarly in both immunosuppressed^6,7^ and immune-competent mice (MMTV-myc^6^ and MMTV-PyMT^19^ mouse models). Although outside of the scope of this manuscript, future plans will involve the study of immune cues regulated by the NR2F1-driven dormancy program in murine melanoma models.

Lastly, MDK can come from cancer cells themselves, but also from stromal fibroblasts, endothelial cells, and infiltrating myeloid cells, particularly under stress, inflammatory, or hypoxic conditions, all contributing to a MDK-rich microenvironment. For instance, aging can induce mammary epithelial cells to secrete MDK to initiate breast cancer formation^20^. Furthermore, in current cutaneous melanoma care, approximately 20–30% of patients who electively discontinue anti-PD-1 therapy eventually relapse, a risk that has been linked to persistent immune activation and chronic inflammation extending beyond the treatment period. Given that MDK is secreted under stress conditions, one might anticipate an increased likelihood of DCC reactivation in these patients. Our findings provide a rationale and mechanism-based strategy to reprogram both the niche and cancer cells to sustain DCC dormancy and prevent metastatic recurrence—through MDK inhibition and NR2F1 activation—as an alternative to traditional cytotoxic approaches or as a means to contain relapse following targeted and/or immune-based therapies.

This research exposes a gap in our understanding of melanoma progression, revealing that certain melanoma DCCs are less prone to becoming dormant before any treatment is applied. Our results show for the first time an interplay between MDK and NR2F1 in lung melanoma DCCs during dormancy.

## Acknowledgements

We extend our sincere gratitude to all the members of the Sosa’s lab including rotation PhD. student Subramanian Thothathri and research trainee Smrithi Jayashree Satheeshkumar. We also acknowledge Ahron Flowers and the Tara Miller Melanoma Center at University of Pennsylvania for the melanoma specimens. We would also like to specially acknowledge Dr. Julio Aguirre-Ghiso for his persistent guidance and the invaluable contributions of his lab members, whose thoughtful discussions during our weekly meetings significantly enriched our research process. We acknowledge Jinghang Zhang from the Flow Cytometry Core and Hillary Guzik at the Analytical Imaging Facility at the Albert Einstein College of Medicine (with support from the NCI Cancer Center Support Grant P30CA013330) for training on the use of The Cytek^®^ Aurora and Hamamatsu S60 slide scanner, respectively. Grant support: MRA Team Science Award N401181 (M.S.Soengas and M.S.Sosa); Gilead Research Scholars (M.S.Sosa.); MRF-CDA (M.S.Sosa).

## Author Contributions

C.R-T. designed, performed in vitro and in vivo experiments, collected microscopy data, analysed data and co-edited the manuscript; T.A.R. performed IF, WB, qRT-PCR experiments, expanded patient-derived melanoma organoids and analysed data; A.A.R. performed 3D, IF, WB, qRT-PCR experiments and analysed data; N.K. IF, WB, qRT-PCR performed experiments and analysed data; A.P-G. performed CDK2 reporter experiments and analysed data; D.O. provided reagents and edited the manuscript; M.S.Soengas provided reagents and edited the manuscript, M.S.Sosa designed and executed CAM assays and in vivo experiments, provided general oversight, collected microscopy data, analysed data and wrote the manuscript.

## Data Availability

The data generated in this study are available upon request from the corresponding author.

## MATERIALS AND METHODS

### Cell lines

Highly metastatic GFP-SK-Mel-147 cells (parental line RRID:CVCL_3876), SK-Mel-28 (RRID:CVCL_0526), WM164 (RRID:CVCL_7928) and B16R2L cells were obtained from Dr. Maria S. Soengas laboratory at CNIO, Spain. SK-Mel-103 (RRID: CVCL_6069) cells were obtained from Dr. Eva Hernando laboratory at NYU, USA. Cells were cultured in DMEM (Dulbecco’s modified medium, Corning 10-031-CV)) supplemented with 10% V/V FBS and 100 U penicillin/ 10 mg/mL streptomycin (Life science # 30-002-CI). Neuroblastoma cell line NB-EBC1 were obtained from Dr. Daniel Weiser at AECOM, US., and cultured in IMDM (Iscove’s Modified Dulbecco’s

Medium, Thermo # 12440053) supplemented with 10% FBS, 1% v/v penicillin/streptomycin, and 1% v/v Insulin-Transferrin-Selenium (ITS, Thermo #41400045).

T-HEp3 cells were derived from a lymph node metastasis from a HNSCC patient as described previously^21^ and kept as PDXs on chick chorioallantoic membrane (CAM). D-HEp3 cells were obtained by passing T-HEp3 cells for more than 40 generations in vitro^21^. In vitro, these cells were cultured in DMEM with 10% of FBS, 100 U/mL penicillin and 0.1 mg/mL streptomycin. All cell lines were tested for mycoplasma contamination every 6 months and used only if tested negative.

### Transfection

D-HEp3 cells were transfected with open reading frame (ORF) lentiviral expression vector pReceiver-Lv105-A0792 (pEX-MDK) and the corresponding empty vector (pEX-Neg) plasmid (obtained from Dr. Maria S. Soengas laboratory at CNIO) by using Fugene6 (Promega # E2693) in serum free and antibiotic-free media according to the manufacturer’s instructions.

Transient knockdown experiments were done using siRNA. Cells were seeded at a density of 20,000 cells/cm^2^ and allowed to reach 60-70% confluency. Cells were then transfected using Lipofectamine^TM^ RNAiMax (Thermo Fisher # 13778150) in serum free media following manufacturer’s instructions. Media was changed 12 hours (h) later and incubated for a total of 48 h before analysis. siRNAs were used at 50 nM unless otherwise specified. For ALK knockdown, cells were transfected with 30 nM ON-TARGETplus Human ALK (238) siRNA - SMARTpool (Horizon # L-003103-00-0005) in serum free and antibiotic-free media according to the manufacturer’s instructions. ON-TARGETplus Non-targeting Control Pool (Horizon # D-001810-10-05) was used as a control.

### Cell cycle analysis

Cell cycle was determined by flow cytometry after EdU incorporation. For this, cells were incubated with 10uM Edu in full media, for 1 h, immediately before collection. Cells were lifted, washed in serum free media and stained with Zombie NIR^TM^ (Biolegend # 423105) for 15 min at room temperature. Cells were washed and fixed on 4% w/v paraformaldehyde (PFA) in PBS for 20 min on ice. Edu was detected by Click-iT^TM^ Edu Alexa Fluor^TM^ 647 kit (Thermo Fisher # C10634) following manufacturer’s instruction. Cells were then stained with DAPI for 15 min on ice (Molecular Probes #P36931). EdU and DNA content was determined on a Cytek Aurora Analyzer at the Flow Cytometry Core Facility at the Albert Einstein College of Medicine. Data was analyzed using FlowJo software (RRID: SCR_008520).

Cell carrying the CDK2-DHB-mVenus reporter (Addgene #136461; RRID:Addgene_136461) were processed for IF. As the CDK2 mediated phosphorylation of this reporter leads to a conformational change that causes the reporter to translocate from the nucleus to the cytoplasm, the ratio of cytoplasmatic to nuclear fluorescence indicates G0/G1 (nuclear only), S (both) or G2/M phases (cytosolic only) were calculated. Images were subjected to cell segmentation using DAPI signal to determine nuclear area and a 2 um expansion to determine cytosolic signal using QuPath software (RRID:SCR_018257). Cell cycle stage was determined by calculating the cytosolic/nuclear ratio of the mean mVenus fluorescence as previously described^15^.

### Lentiviral transduction

293T cells (RRID: CVCL_0063) were seeded at 35,000 cells/cm^2^ in full DMEM media. Once cultures reached 80% confluency, they were transfected with 2.5 μg of lentiviral vectors using Lenti-pack HIV Expression Packaging Kit (GeneCopoeia # LT001) in serum free media. Cell culture media was replaced with full media supplemented with Titerboost^TM^ reagent (1:500 dilution) 12 hours after transfection. Supernatants enriched in viral particles were collected 48 h and 72 h later. Pooled supernatants were filtered through a 0.45-μm filter and concentrated using Lenti-X concentrator (Takara # 631231). Aliquots were used immediately or stored at -80°C for 6 months.

To generate stable cell lines, SK-Mel-147 and SK-Mel-28 cells were transduced with lentiviral particles carrying CDK-DHB-mVenus^15^, shControl or shMDK constructs in full media supplemented with 0.8 μg/mL Polybrene (EMD Millipore # TR-1003-G). Cell culture media was replaced 24 h after, and selection of transduced cells was carried by treatment with 1ug/ml puromycin 72 h post-infection. Targeting/non targeting sequences for shRNA against MDK were obtained from Dr. Maria S. Soengas (see Table 3).

**Table 1:**
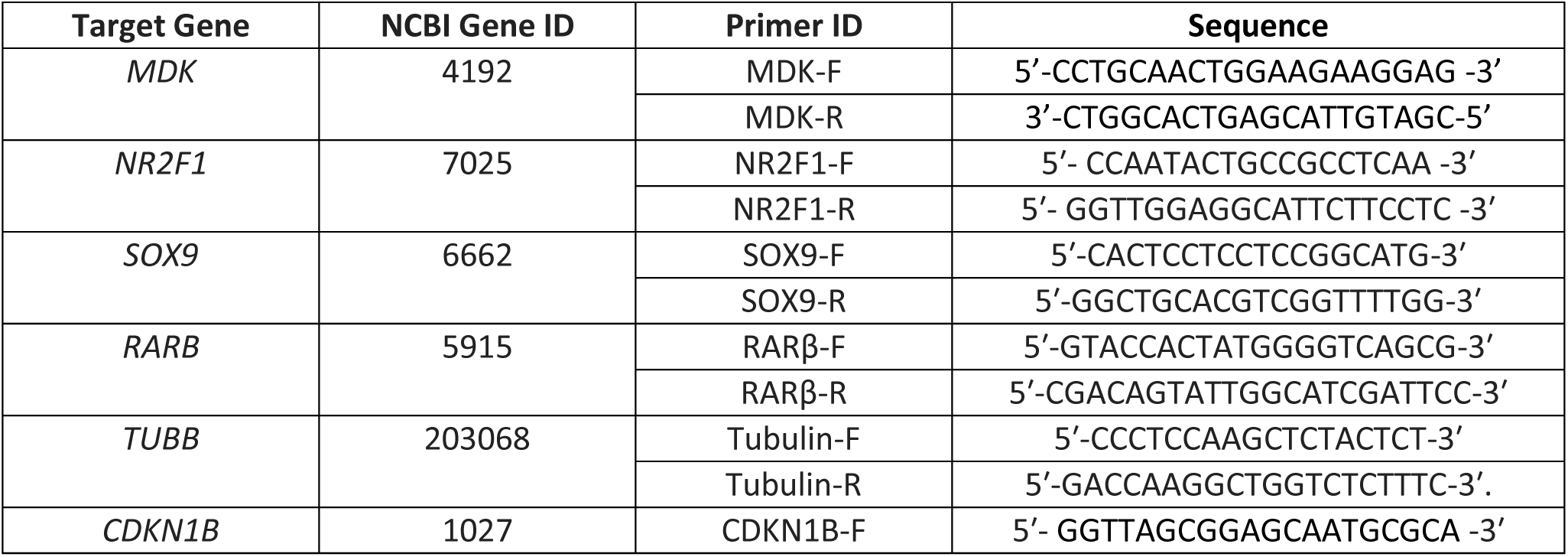
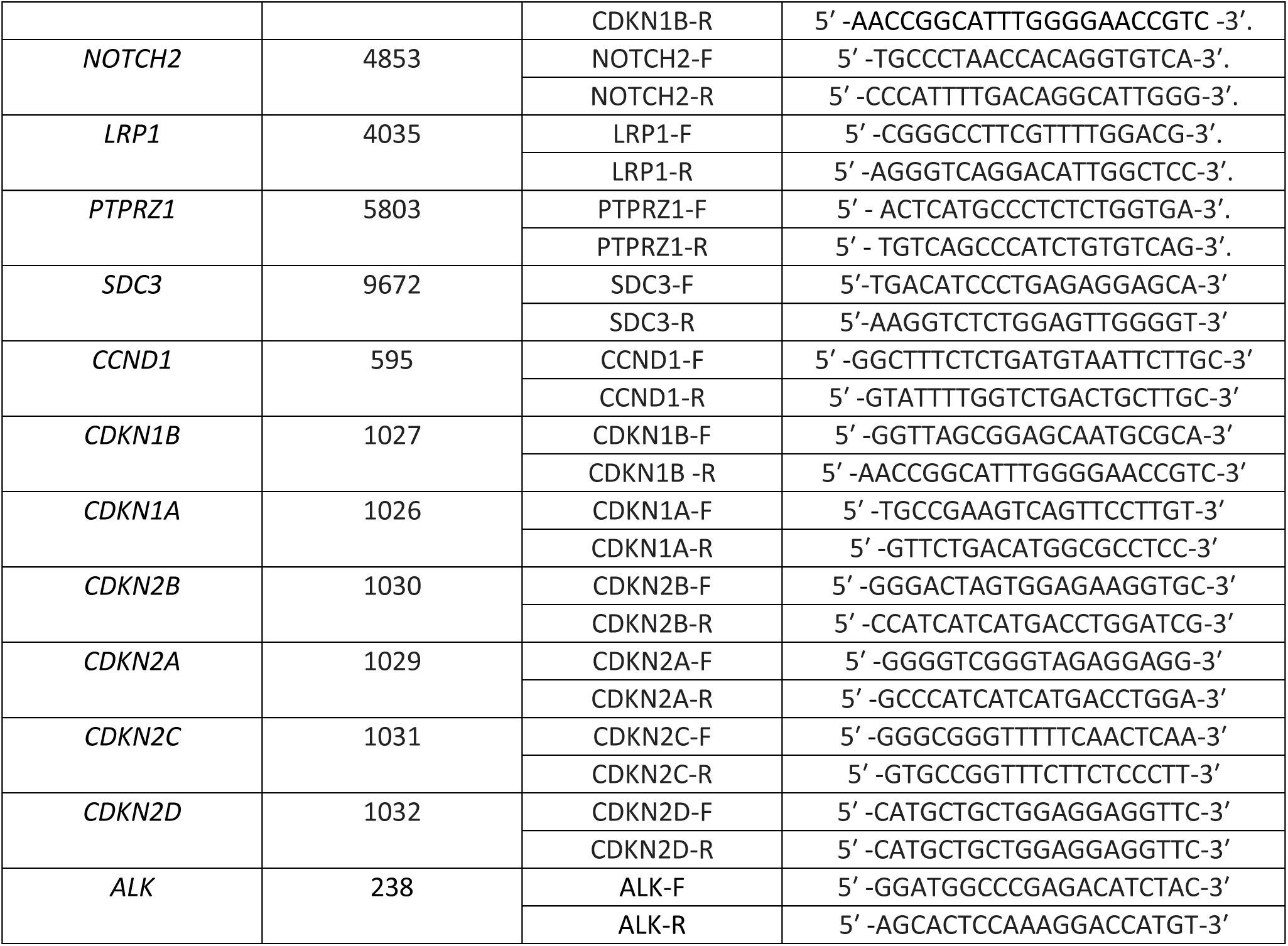
qRT-PCR primers.

**Table 2:**
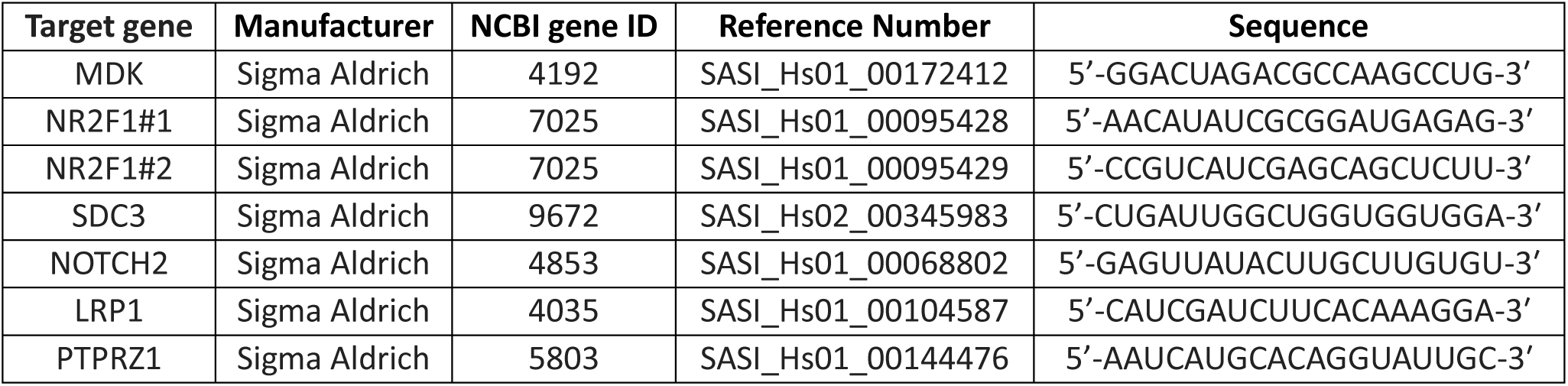
siRNA targeting sequences.

**Table 3:**
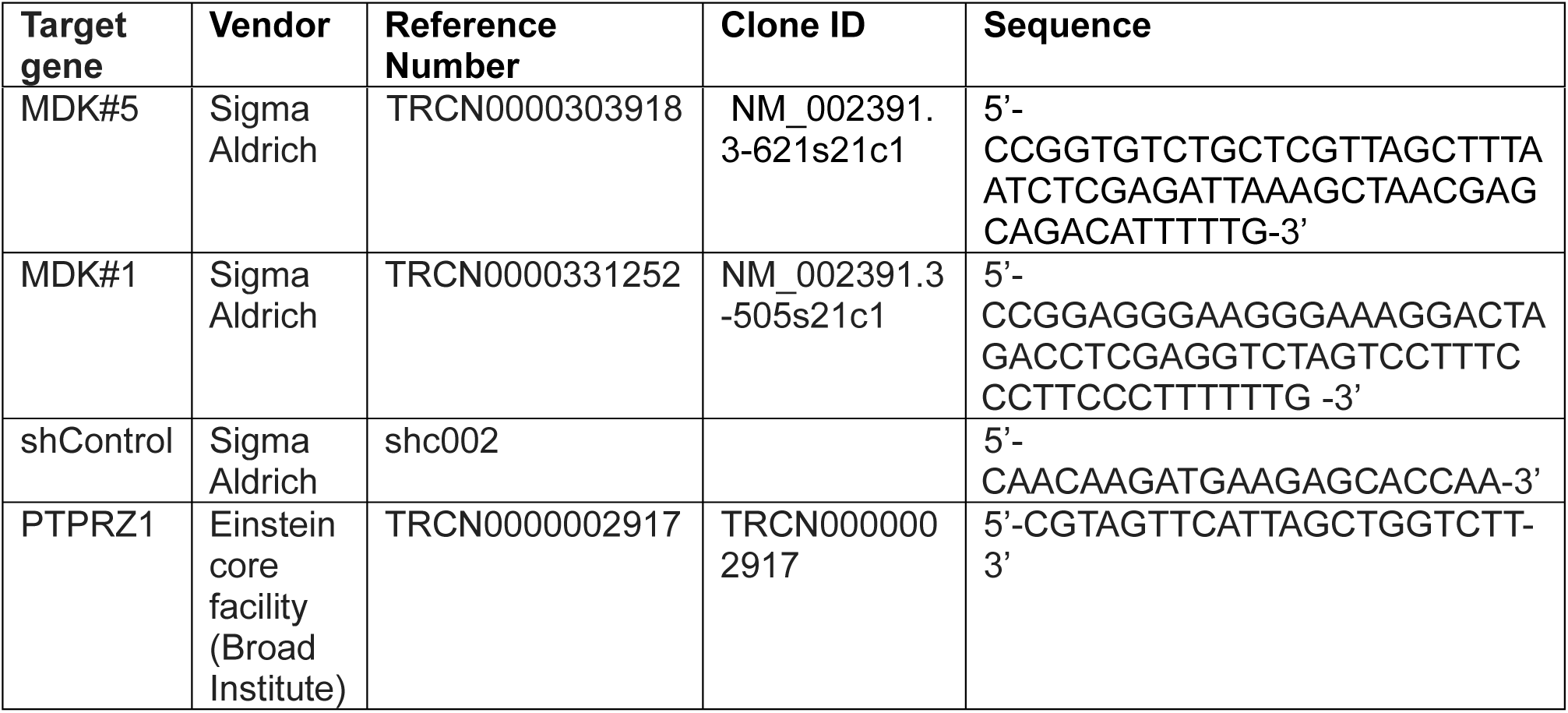
shRNA targeting sequences.

**Table 4:** Antibody list

| Antibody Name | Manufacturer | Catalog Number | RRID | Use |
| --- | --- | --- | --- | --- |
| NR2F1 | Abcam | 181137 | AB_2890250 | WB (1:300)<br>IF (1:100)<br>MICSSS (1:100) |
| SOX9 | Millipore | AB5535 | AB_2239761 |  |
| CDKN1B | CST | 3686 | AB_2077850 | IF (1:100) |
| phospho-p38 | CST | 9211S | AB_331641 | WB (1:1000) |
| p38 | CST | 9212 | AB_330713 | WB (1:1000) |
| phospho-ERK 1/2 | CST | 9106 | AB_331768 | WB (1:1000) |
| ERK 1/2 | CST | 9102 | AB_330744 | WB (1:1000) |
| B-Tubulin | Abcam | Ab15568 | AB_2210952 | WB (1:1000) |
| GAPDH | CST | 2118 | AB_561053 | WB (1:1000) |
| Lamin B1 | CST | 12586S | AB_2650517 | WB (1:500) |
| phospho-AKT | CST | 4060s | AB_2315049 | WB (2:1000) |
| AKT | CST | 4685s | AB_2225340 | WB (1:1000) |
| phospho-ALK | CST | 3341s | AB_331047 | WB (2:1000) |
| phospho-ERK | CST | 4377s | AB_331775 | WB (1:1000) |
| ALK | CST | 3633 | AB_11127207 | WB (1:1000) |
| MK-9 (MDK) | Santa Cruz | sc-46701 | AB_627949 | IF (1:50)<br>MICSSS (1:50) |
| HMB45 | Santa Cruz | sc-59305 | AB_784471 | IF (1:50)<br>MICSSS (1:100) |
| Vimentin | R&D | MAB21051 | AB_2216114 | IF (1:100) |
| ATF2 | CST | 35031 | AB_2799069 | WB (1:1000) |
| Phospho-ATF2 | Abcam | ab32019 | AB_725567 | WB (1:500) |
| MK2 (MAPKAPK2) | CST | 3042 | AB_10694238 | WB (1:1000) |
| p-MK2 | CST | 3041 | AB_330726 | WB (1:1000) |
| RARB | Abcam | ab15515 | AB_301924 | WB (1:1000) |
| Histone H3 | CST | 9717 | AB_331222 | WB (1:1000) |
| β-Actin | CST | 3700 | AB_2242334 | WB (1:1000) |
| Phospho-S6 | CST | 5364 | AB_10694233 | WB (2:5000)<br>IF (1:150) |
| S6 | CST | 2217 | AB_331355 | WB (2:5000) |
| AF488 anti-mouse | Thermo | A-11001 | AB_2534069 | IF (1:1000) |
| Phosphor-AMPKα | CST | 2535 | AB_331250 | WB (1:1000) |
| AMPKα | CST | 2532 | AB_330331 | WB(1:1000) |
| AF568 anti- rabbit | Thermo | A-11036 | AB_10563566 | IF (1:1000) |
| Ki-67-APC | Miltenyi | 130-120-556 | AB_2784389 | IF (1:150) |
| AF488 anti-goat | Thermo | A11055 | AB_2534102 | IF (1:1000) |
| AF647 anti-rabbit | Thermo | A-31573 | AB_2536183 | IF (1:1000) |

### CAM assays

Tumor growth of T-HEp3 cells on CAM has been already described^22^. Briefly, Control or ORF-MDK D-HEp3 cells were inoculated on CAM and allowed to grow for 7 days. Tumors were harvested and digested with collagenase 1A (Sigma-Aldrich; C9891) for 30 min at 37°C. D-HEp3 tumor cells (recognized by their very large size compared with chicken cells) were counted using a hemacytometer.

### Animal models

Animal procedures were approved by the Institutional Animal Care and Use Committee (IACUC) of Icahn School of Medicine at Mount Sinai with protocol 2017-0162, and the Albert Einstein College of Medicine with protocol 00001676. Nude mice were purchased from Jackson Laboratories (Stock #002019; RRID:IMSR_JAX:002019) and maintained under specific pathogen free (SPF) conditions throughout the experiment.

To determine spontaneous metastatic burden, 6 weeks old nude females were subcutaneously injected with melanoma cells. Cells had been expanded, collected at 80% confluency, and single cells resuspended in 1X PBS. Animals were randomized into two groups and injected with 100 uL of cell suspensions in a 1:1 PBS:Matrigel solution (1,000,000 cells/mouse) using a 27G needle. Mice were euthanized, and lungs were collected after perfusion. Animals were maintained under SPF conditions throughout the experiment and monitored twice weekly.

To determine perform experimental metastasis assays- in absence of primary tumors-, 6 weeks old nude females were intravenously injected with melanoma cells. Cells had been expanded, collected at 80% confluency, and single cells resuspended in 1X PBS. Animals were randomized into two groups and injected with 100 uL of cell suspensions (1,000,000 cells/mouse) on the tail vein using a 27G needle. Twenty-one days later, mice were euthanized, and lungs were collected after perfusion. Animals were maintained under SPF conditions throughout the experiment and monitored twice weekly.

To determine the effect of C26 (Chembridge #sc-6596020) on metastatic burden, nude female animals of 6 weeks of age were intravenously injected with shControl or shMDK cells (1x10^6^ cells/mouse in 100uL) on the tail vein using a 27G needle. Three days after injection, animals were randomized into groups receiving vehicle (30% v/v polyethylene glycol 300, 5% v/v polysorbate 80 and 4% v/v dimethyl sulfoxide (DMSO) in sterile water) or treatment with C26 (0.5mg/Kg) by intraperitoneal injection every other day, for 28 days. Animals were maintained under SPF conditions throughout the experiment and monitored thrice weekly. Twenty-eight days later, mice were euthanized, and lungs were collected after perfusion.

### Patient Samples

Paraffin embedded sections from patient melanoma tumors and lymph nodes (positive and negative for metastasis by histopathological analysis) were obtained from Dr. Lynn Schuchter from the University of Pennsylvania (MTA ID: 44613/00, MTA ID:000344-2025).

### 3D cultures

WM164 cells were grown in 3D cultures in 8-well chambers (Corning # 354118). For this, 8 well chambers were coated with a layer of growth factor reduced Matrigel® (Corning # 354230). After Matrigel® was solid, 1,000 cells were seeded on top, in complete media and allowed to form colonies for 48 h. Then, cells were treated with vehicle or 20 ng/mL recombinant human Midkine rhMDK for 48 h. Brightfield images were taken on Leica DMi8 scope (20X, DIC) daily. On day 5, cells were fixed using 4% w/v PFA for 20 min at room temperature and process for immunofluorescence.

SK-Mel 147 shControl cells pretreated with 1uM Compound 26^7^ and 25nM iMDK (3-[2-[(4-Fluorophenyl)methyl]imidazo[2,1-b]thiazol-6-yl]-2H-1-benzopyran-2-one; EMD Millipore # 508052) for 48 h in 2D. Cells were then seeded in 8 well chambers (1000 cells/well) over a layer of Matrigel and maintained treatment with C26, iMDK, or a combination of both drugs for 6 days. Brightfield images were taken daily on a Zeiss Axio Observer scope (20X, differential interference contrast (DIC)).

### Western blot

Whole cell lysates were obtained from SK-Mel-147 shControl and shMDK cells grown at 70-80% confluency and serum-starved for 20 h. Samples were collected in 1xRIPA buffer and centrifuged at 12000rpm and 4°C to clarify lysates. Cell fractionated lysates were obtained from cells seeded at a density of 36,000 cells/cm2 and treated with P38 inhibitor SB203580 (Sellechem # S1076) or MDK inhibitor iMDK (EMD Millipore # 508052) in serum-free medium for 24 hours. To test the responses to Alecitinib treatment, SK-Mel-147 parental cells were treated with 0.5 uM or 1 uM Alecitinib in full media for 24 h. Cells were washed once with cold 1X-PBS and processed for nuclear and cytosolic protein fractionation using a cytoplasmic and nuclear protein extraction kit (Boster bio # AR0106). Protein concentration was determined using BCA protein assay kit (Pierce #23227) and a standard bovine serum albumin (BSA) curve. Samples were then boiled for 5 min at 95°C in SDS-Page sample loading buffer (Bioscience # 786-701). SDS-PAGE gels at 8 or 10% were run in Running Buffer (25mM Tris, 190mM glycine, 0.1% w/v SDS) and transferred to polyvinylidene fluoride (PVDF) membranes in Transfer Buffer (25mM Tris, 190mM glycine, 20% v/v methanol). Membranes were then blocked in 5% w/v milk in TBST (Polysorbate 20-TBS buffer; 50mM Tris, 150mM Sodium Chloride, 0.1% Polysorbate 20, pH7.6). Membranes were incubated in primary antibodies at 4°C overnight. Following 3 washes with TBST buffer, membranes were incubated with horseradish peroxidase (HRP)-conjugated secondary antibodies at room temperature for one hour. Western blot development was done using Amersham ECL Western Blot Detection (GE, RPN 2106) and GE ImageQuant LAS 4010.

### Immunofluorescence staining

Cells in two-dimensional (2D) cultures (SK-Mel 147 shC, shMDK and Wm164 cells) were seeded on 12mm coverslip and, after 24 h, fixed with 4% w/v PFA. Cells were permeabilized using 0.1% v/v Triton X-100 in PBS for 20min, treated with 50mM NH4Cl, and blocked using 3% BSA in PBS + 10% Normal Goat Serum (Gibco #PCN5000) for one hour. Coverslips were then incubated with primary antibodies for MDK and NR2F1 overnight at 4C. Secondary antibodies were used at 1:1000 dilution in 3% w/v BSA in PBS. Coverslips were mounted with ProLong™ Gold Antifade reagent with DAPI (Invitrogen #P36931).

Lymph node and primary tumors sections from 10 patients were stained using multiplexed Immunohistochemical consecutive staining in a single slide (MICSSS)^23^ using antibodies against human Midkine, NR2F1 and HMB45 (see antibody list). For this, sections were baked overnight at 60°C and deparaffinized and rehydrated by consecutive xylene and ethanol washes. Heat-induced antigen retrieval was done for 30 min at pH6 with a 10mM citrate buffer. Slices were rinsed with TBS (10mM Trizma, 150mM NaCl, pH7.4) and permeabilized at room temperature (RT) for 5 min in 0.1% v/v Triton X100 in TBS. Endogenous peroxidase was quenched by incubation in 3% H2O2 for 15 min at RT before blocking with SuperBlock (Thermo Fisher #37515) for 30 min. Tissues were then incubated with an anti-HMB45 antibody diluted in TBS at 4°C overnight, washed in 0.04% v/v Polysorbate 20 in TBS for 5 min and incubated with HRP-conjugated secondary antibodies at 1:500 dilution in TBS for 1 h at room temperature. Slides were developed using aminoethyl carbazole (AEC) solution following manufacturer’s instruction (Vector # SK-4200, RRID:AB_2336076), counterstained with Harris hematoxylin, and mounted in Glycergel (Dako # C056330). Sections were imaged using the Hamamatsu Slide Scanner S60 at a 20X magnification. Then, sections were subjected to consecutive rounds of demounting, destaining, re-staining and imaging. For demounting, sections were incubated in warm water (55°C) until coverslips slipped off on its own and washed in water. For destaining, sections were immersed in 50% v/v ethanol for 2 min, 1.2N HCL in 70% v/v ethanol for 2 min, 100% v/v ethanol for 5 min, 70% v/v ethanol for 2 min, 50% v/v ethanol for 2 min, and water for 5 min. Slides were monitored at this step to confirm destaining of AEC and hematoxylin by microscopy. Slides were then subjected to a second round of antigen retrieval for 10 min at pH6 with a 10mM citrate buffer or at pH9 using a Tris-based solution (Dako # H-3301, RRID:AB_2336227), depending on primary antibody. Blocking with Fab fragments (Jackson ImmunoResearch #715-007, RRID:AB_2307338; #711-007, RRID:AB_2340587; or #712-007, RRID:AB_2340634) in TBS was followed by primary and secondary antibody staining, development, mounting and imaging. As a last round, sections were processed for fluorescent staining with antibodies against HMB45, phosphoS6 (pS6) and Ki-67 (see antibody list), using Hoechst 33342 (Thermo Fisher #H3570, RRID:AB_3675235) as counterstaining. All images were acquired using a Hamamatsu S60 Nanozoomer scanner (Analytical Imaging Facility, Albert Einstein College of Medicine), stacked and aligned using the Linear stack alignment with SIFT plugin in FIJI (ImageJ, RRID: SCR_003070). HMB45+ regions were annotated using QuPath and classified as DCCs (single cell), small clusters (between 2 and 10 cells), micromets (between 11 and 100 cells) and macromets (above 100 cells). These regions were then scored based on MDK and NR2F1 staining: first as negative or positive based on the presence of specific cytoplasmic and/or membranous signal (for MDK) or nuclear signal (for NR2F1) in tumor cells. Among positive cases, expression was further categorized as ‘low’ or ‘high’ according to the proportion of stained tumor cells and staining intensity: low’ represents weak to moderate staining in up to 50 percent of tumor cells; ‘high’ represents strong staining in more than 50 percent of tumor cells. Cell segmentation over Hoechst 33342 signal (nuclear area) and cell expansion of 2um (cytosol) were used to determine the mean fluorescence signal of Ki-67 and pS6, respectively.

Mouse tissue sections were deparaffinized and rehydrated as before. Antigen retrieval was done in 10 mM sodium citrate buffer (pH 6) and permeabilized in 0.1% v/v Triton X-100 in PBS. Blocking was done using 3% w/v BSA in PBS with 1.5% v/v normal goat (MP Biomedicals #8642921) or normal donkey serum (Sigma-Aldrich #D9663; RRID:AB_2810235). Primary antibodies at 1:100 dilution unless otherwise specified–were used overnight at 4°C. Secondary antibodies at 1:1000 dilution were assessed for 1 hour at room temperature. Slides were mounted using Prolong Gold Antifade mounting media with DAPI (Molecular Probes #P36931). Image acquisition was done using Leica DMi8 fluorescence microscope and image processing was done using LAS X Leica software (RRID:SCR_013673) and FIJI software.

### Flow Cytometry

For pharmacological inhibition of ALK, SK-Mel-147 and NB-EBC1 cells were seeded in 2D cultures in low serum media (2-5% v/v) overnight, treated with 1uM Alectinib (Selleckchem #S5232) or 25nM Zotizalkib (Selleckchem #S9959) for 1 h at 37C, and stimulated with recombinant MDK for 15 min. Cells were collected in presence of phosphatase inhibitors (5mM Sodium Fluoride and 5mM beta-glycerolphosphate) by trypsin treatment and fixation in 4% w/v in PFA. Cell suspensions were permeabilized in 90% v/v methanol for 1 h on ice and stained with primary antibodies against ALK and phospho-ALK (pY1586) for 1h on ice. Secondary anti-rabbit antibodies conjugated to Alexa-Fluor 488 were used. Negative gating was done based on secondary antibody controls (no primary). NB-EBC1 cells were used as positive controls for ALK expression.

### Quantitative reverse transcription PCR (qRT-PCR)

RNA was extracted from 2D and 3D cell cultures using Trizol^TM^ (Thermo Fisher # 15596026) or using Qiagen Micro RNeasy columns (Qiagen #74004). RNA (1-2 ug) was treated with DNase I (Thermo Fisher # AM2222) and retrotranscribed using RevertAid First Strand cDNA Synthesis Kit (Thermo Fisher #K1622). Quantitative real time-PCR was performed using PerfeCTa SYBR® green FastMix(R) Reaction mix (Quantabio #95072) in Agilent QuantStudio3 (RRID:SCR_018712). *TUBULIN (TUBB)* was used as housekeeping control for all plates and gene expression determined using the ΔΔCt method.

### Statistical Analysis

Statistical analysis was done using Prism Software (RRID:SCR_002798). Differences were considered significant if *P* values were <0.05. For the majority of cell culture experiments, unless specified otherwise, one tailed unpaired *t-test* was used. For mouse experiments, one tailed unpaired *Mann-Whitney* tests was used. Chosen sample size was determined empirically.

Reagents lists

## Notes

### Competing Interest Statement

The authors have declared no competing interest.

### Summary of Updates

The manuscript is updated to accommodate new findings.

