## Supplemental Figures for "Midkine Drives Reawakening of Dormant Melanoma Metastases"

**Running Title:** Midkine inhibits dormancy.

**#** equal contributions.

***Materials and Correspondence:*** Maria Soledad Sosa, Department of Microbiology and Immunology, Albert Einstein College of Medicine, Bronx, NY 10461, USA. Phone: (+1) 718.430.2115.

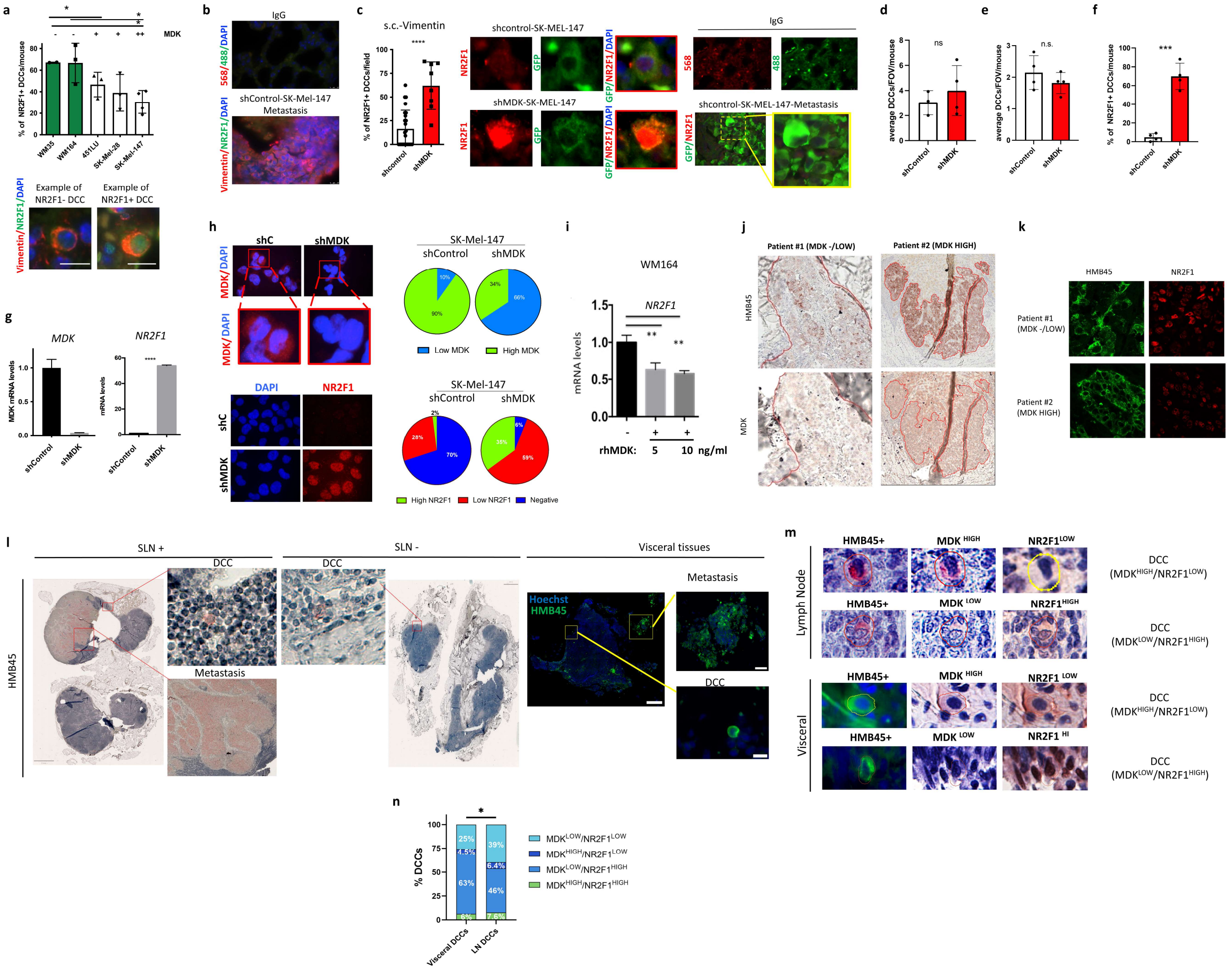


**Supplementary Figure 1. Midkine represses NR2F1 levels in melanoma**. **a.** WM35, WM164, 451LU, SK-Mel-28 and SK-Mel-147 melanoma cells were s.c. injected into nude mice. Lung sections were stained for NR2F1 (green), Vimentin (red) and DAPI (blue). Bar plots show percentage of NR2F1+ DCCs/mouse. Unpaired t-test. Images show examples of single NR2F1- and NR2F1+ DCCs. Scale bars represent 10 μm. **b.** Pictures show IgG (upper panel) and staining for NR2F1 (green), Vimentin (red), DAPI (blue) in a metastasis found in shControl SK-Mel-147 s.c. injected mice (lower panel). Scale bars represent 10 μm. **c.** GFP-SK-Mel-147 (shControl and shMDK) cells were implanted s.c. in nude mice. Lungs were collected 21 days later and embedded in paraffin. Lung sections were stained for NR2F1 (red), GFP (green) and DAPI (blue). Percentage of NR2F1+ DCCs/field in shControl- or shMDK-SK-MEL-147 lungs. Mann-Whitney test. **d.** Average number of DCCs/field of view (FOV)/mouse was plotted from same animals as in **Fig. 1b**. Unpaired t-test. **e.** SK-Mel-28 (shControl and shMDK) cells were implanted s.c. in nude mice. Lungs were collected 21 days later and embedded in paraffin. Lung sections were stained for NR2F1 (red), GFP (green) and DAPI (blue) as in **Fig. 1b**. Average number of DCCs/field of view (FOV)/mouse was plotted. Unpaired t-test. **f.** Percentage of NR2F1+ DCCs/mouse in shControl- or shMDK-SK-MEL-28 lungs was plotted. Unpaired t-test. **g.** *MDK* and *NR2F1* mRNA levels in shControl- and shMDK SK-Mel-147 cells. **h.** shControl and shMDK SK-Mel-147 cells were seed in 2D and stained for NR2F1 and MDK by IF. Representative images for NR2F1 and MDK are shown per cell line. Pie graphs show percentages for the indicated categories over total number of cells quantified. **i.** WM164 cells were treated with rhMDK at the indicated concentrations for 48 h. Levels of *NR2F1* were measured by qRT-PCR. Unpaired t-test. **j&k. j**: Primary tumors from melanoma patients were stained for HMB45 and scored for MDK based on IHC staining. **k**: Same tumors as in **j** were stained by IF for NR2F1 (red) and HMB45 (green). Quantification of the percentages of NR2F1+ tumor cells (nuclear staining) is plotted in **Fig. 1d. l.** LN-, LN+ and visceral (brain, lungs and omentum) samples were sequentially stained by MICSSS for the indicated antigens. Quantification plots were shown in **Fig. 1e**. **m.** Representative images of a single DCC (positive for HMB45) with $MDK^{HIGH}/NR2F1^{LOW}$ or $MDK^{LOW}/NR2F1^{HIGH}$ phenotype from LN and visceral biopsies. **n.** Dormant DCCs (Ki-67 negative) from visceral (N=244) and LN (N=78) biopsies were identified and the percentages of DCCs with the indicated phenotypes were plotted. Fisher exact's test (p value <0.05).

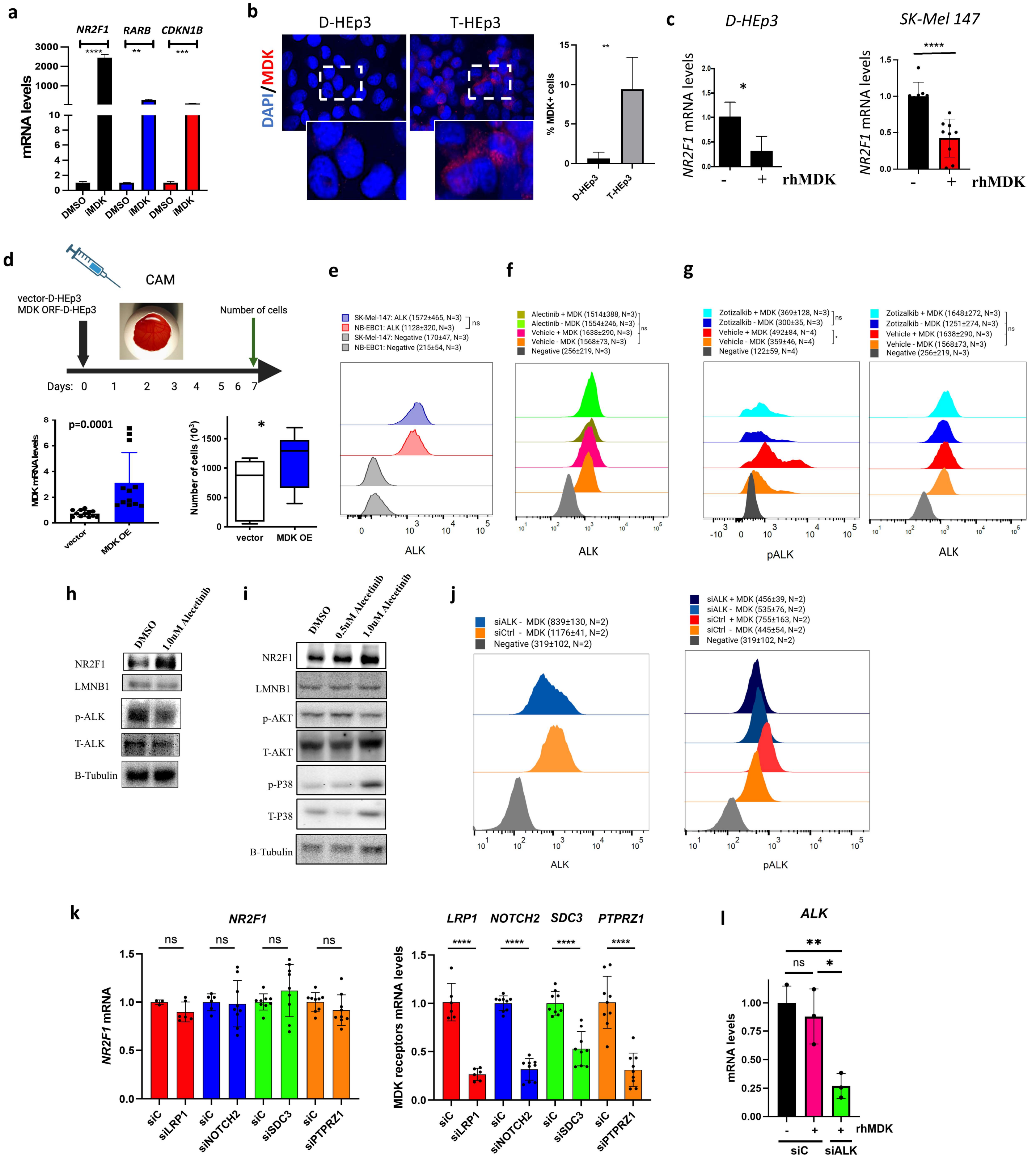
a
NR2F1
RARB
CDKN1B
mRNA levels
DMSO
iMDK
b
D-HEp3
T-HEp3
DAPI/MDK
% MDK+ cells
c
D-HEp3
SK-Mel 147
NR2F1 mRNA levels
rhMDK
d
CAM
vector-D-HEp3
MDK ORF-D-HEp3
Number of cells
Days:
p=0.0001
MDK mRNA levels
vector
MDK OE
Number of cells ($10^3$)
e
SK-Mel-147: ALK (1572±465, N=3)
NB-EBC1: ALK (1128±320, N=3)
SK-Mel-147: Negative (170±47, N=3)
NB-EBC1: Negative (215±54, N=3)
ALK
f
Alectinib + MDK (1514±388, N=3)
Alectinib - MDK (1554±246, N=3)
Vehicle + MDK (1638±290, N=3)
Vehicle - MDK (1568±73, N=3)
Negative (256±219, N=3)
ALK
g
Zotizalkib + MDK (369±128, N=3)
Zotizalkib - MDK (300±35, N=3)
Vehicle + MDK (492±84, N=4)
Vehicle - MDK (359±46, N=4)
Negative (122±59, N=4)
pALK
Zotizalkib + MDK (1648±272, N=3)
Zotizalkib - MDK (1251±274, N=3)
Vehicle + MDK (1638±290, N=3)
Vehicle - MDK (1568±73, N=3)
Negative (256±219, N=3)
ALK
h
DMSO
1.0uM Alecetinib
NR2F1
LMNB1
p-ALK
T-ALK
B-Tubulin
i
DMSO
0.5uM Alecetinib
1.0uM Alecetinib
NR2F1
LMNB1
p-AKT
T-AKT
p-P38
T-P38
B-Tubulin
j
siALK - MDK (839±130, N=2)
siCtrl - MDK (1176±41, N=2)
Negative (319±102, N=2)
ALK
siALK + MDK (456±39, N=2)
siALK - MDK (535±76, N=2)
siCtrl + MDK (755±163, N=2)
siCtrl - MDK (445±54, N=2)
Negative (319±102, N=2)
pALK
k
NR2F1
NR2F1 mRNA
siC
siLRP1
siNOTCH2
siSDC3
siPTPRZ1
LRP1
NOTCH2
SDC3
PTPRZ1
MDK receptors mRNA levels
l
ALK
mRNA levels
rhMDK
siC
siALK

**Supplementary Figure 2. Midkine overexpression favors reactivation of dormant cells. a.** SK-Mel-147 cells were treated with MDK inhibitor (MDKi, 25nM) for 48 h and qRT-PCR analysis for the indicated genes were performed. Unpaired t-test. **b.** D-HEp3 and T-HEp3 cells were seeded in 2D and stained for MDK (red) by IF. Percentage of MDK positive cells in each cell line is plotted. Representative images are shown. **c.** D-HEp3 (left) and SK-Mel 147 (right) cells were daily treated with rhMDK for 48 h and qRT-PCR levels for *NR2F1* were measured. **d.** D-HEp3 cells were transfected with empty pEX or pEX-MDK. Next days, cells were inoculated into chicken embryos (5 eggs/condition). Five days later, absolute number of tumor cells were counted. The level of *MDK* mRNA were measured by qRT-PCR. Unpaired t-test. **e**. Untreated SK-Mel-147 cell and NB-EBC1 cells (+ control) were fixed and stained with anti-ALK antibodies by flow cytometry. Histograms show the MFI for each line and its respective controls. N=3. MFI+/- SD is shown. 2-way ANNOVA. **f**. Total ALK level in SK-Mel-147 treated with alectinib for 1h, and rhMDK for 15 min, determined by flow cytometry. N=3. 2-way ANNOVA. **g**. Total ALK and pALK in SK-Mel-147 cells seeded in 2D, treated with 25nM Zotizalkib for 1h, and 20ng/mL rhMDK for 15 min. Determined by flow cytometry. Histograms show MFI. N=3-4. 2-way ANNOVA (* p-value <0.05). **h**. SK-Mel-147 cells were seeded in 2D and then treated with Alectinib for 24 hours. Western blots for the indicated targets are shown. Nuclear extraction was used to detect NR2F1 and LAMIN B1. **i.** B16R 2L cells were treated with Alectinib (0.5 μM and 1.0μM) for 24 h and the target proteins were checked by immunoblotting analysis. Nuclear extraction was used to detect NR2F1 and LAMIN B1. **j.** Total and phospho-ALK determined by flow cytometry in SK-Mel-147 cells seeded in 2D, transfected with siControl and siALK for 48 hours and stimulated with 20ng/mL rhMDK for 15 min. Histograms show MFI. N=2. MFI+/-SD is shown. **k**. SK-Mel-147 cells were seeded in 2D and then transfected with siControl and siRNAs against the indicated receptors for 48 hours. qRT-PCR for the indicated targets is shown. Results from two or three independent experiments. Unpaired t-test. **l**. SK-Mel 147 cells were daily treated with rhMDK for 72 h with or without siALK and qRT-PCR levels for *ALK* were measured. Unpaired t-test.

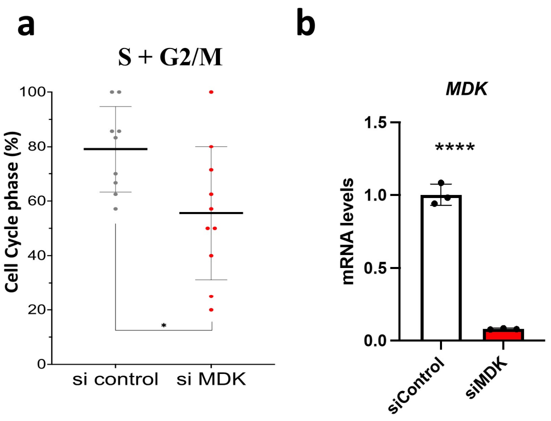


**Supplementary Figure 3. Midkine depletion induces cell cycle arrest. a.** SK-Mel-147 cells stably expressing CDK2-mVenus reporter were transfected with siControl or siMDK.After 48 hours, cells were fixed and the percentage of cells in S+G2/M phase was plotted. Unpaired t-test. **b.** SK-Mel-147 cells were transfected with siMDK or siControl. Graph shows the efficacy of *MDK* knockdown. Unpaired t-test.

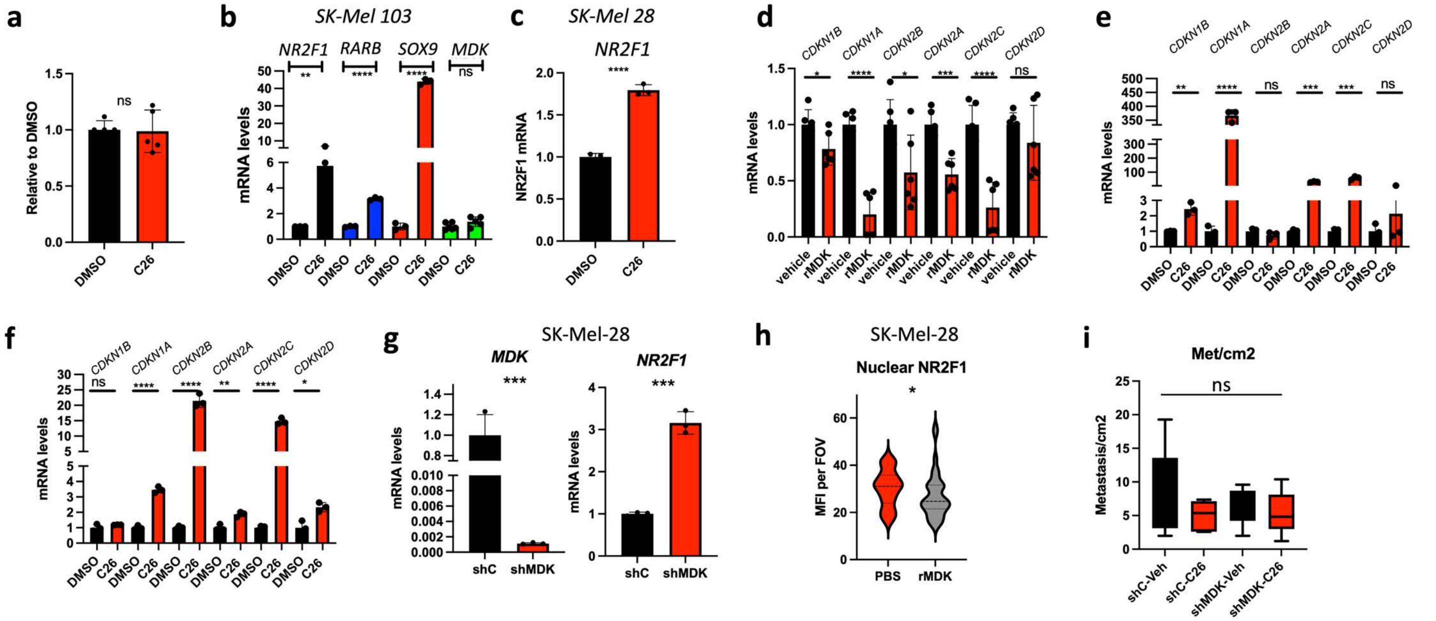


**Supplementary Figure 4. NR2F1 agonist induces NR2F1 downstream genes while addition of recombinant MDK antagonizes this effect. a.** Relative secreted MDK levels measured by ELISA after C26 (1 μM) in SK-Mel-147 cells. Unpaired t-test. **b**. Relative mRNA levels of *NR2F1, RARB, SOX9 and CDKN1B* in DMSO and C26 treated SK-Mel- 103 cells. Unpaired t-test. **c.** SK-Mel-28 cells were treated with C26 for 24 h and NR2F1 levels were measured by qRT-PCR. Unpaired t-test. **d.** SK-Mel-147 cells were treated with rhMDK (20ng/ml) for 24 h and the indicated genes were measured by qRT-PCR. Unpaired t-test. **e.** SK-Mel-147 cells were treated with C26 for 24 h and the indicated genes were measured by qRT-PCR. Unpaired t-test. **f.** SK-Mel-28 cells were treated with C26 for 24 h and the indicated genes were measured by qRT-PCR. Unpaired t-test. **g**. Relative mRNA levels of *NR2F1* in shControl SK-Mel-28 cells vs. shMDK-SK-Mel-28 cells. Unpaired t-test. **h.** Nuclear NR2F1 levels (MFI, mean fluorescence intensity) in PBS vs. rhMDK-treated SK-Mel-28 cells. **i**. shControl and shMDK SK-Mel-28 cells were i.v. injected (one million cells/mouse) into nude mice followed by treatment with vehicle or C26 for one month. Collected lung were stained by H&E and number of metastasis were measured. N=5 mice/group. Mann Whitney.

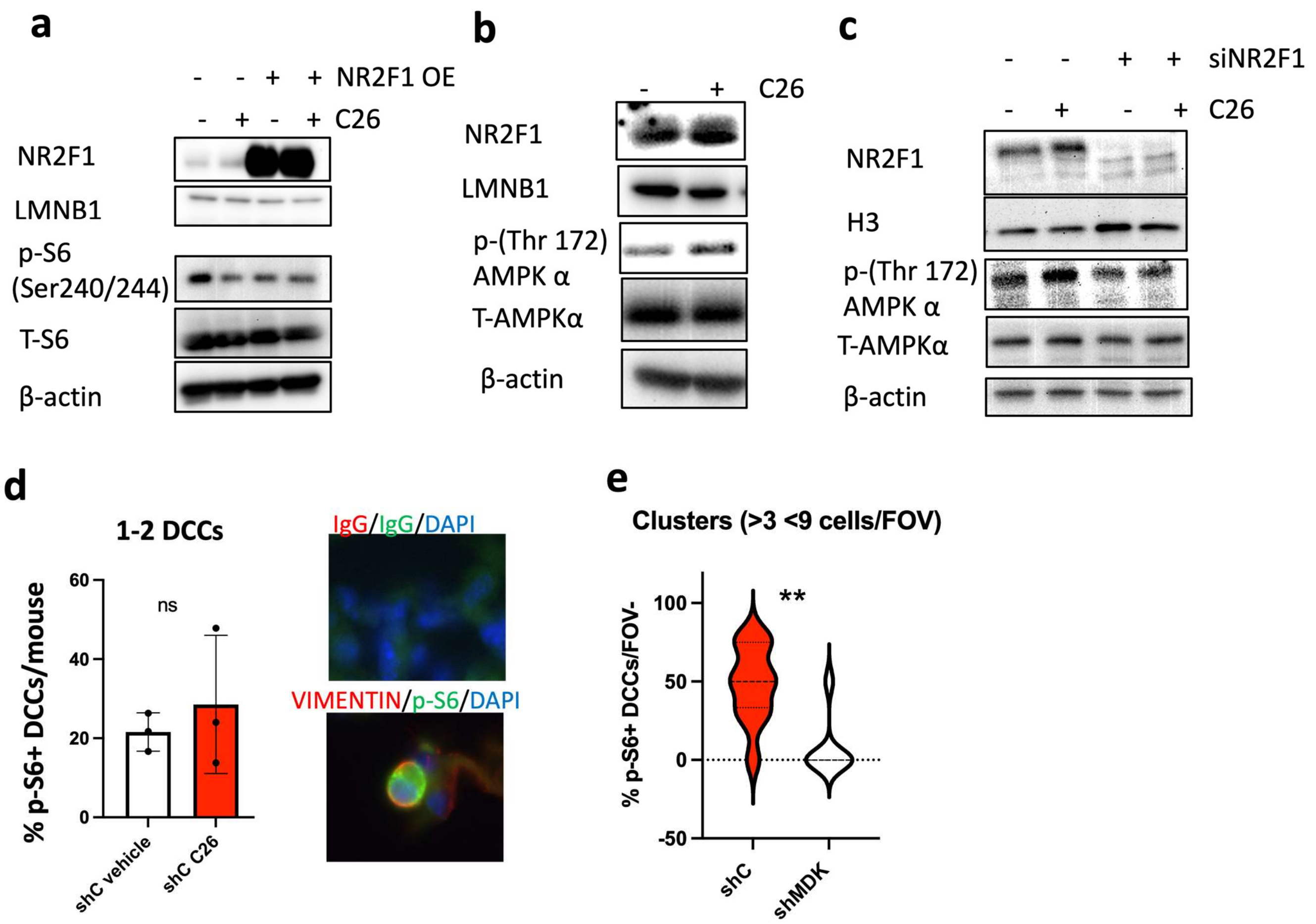


**Supplementary Figure 5. NR2F1 activation blocks mTOR signaling in MDK$^{HIGH}$ melanoma. a**. SK-Mel-147 cells were transfected with NR2F1 plasmid or empty vector and treated with C26 or vehicle for 24 h and nuclear and cytosolic fractions were collected and plotted against the indicated antigens. **b.** SK-Mel-147 cells were treated with C26 for 24 h and nuclear and cytosolic fractions were collected and plotted against the indicated antigens**. c.** SK-Mel-147 cells were depleted of NR2F1 and treated with C26 for 24 h and nuclear and cytosolic fractions were collected and plotted against the indicated antigens**. d.** Collected lungs shown in **Fig. 4c** were stained by IF for Vimentin and PS6 and the percentage of p-S6+ DCCs was plotted (N= 104 DCCs in shC, N= 105 DCCs in shC C26). Representative image of a positive p-S6 DCC (bottom) and its IgG control (top). **e**. Collected lungs shown in **Fig. 4c** were stained by IF for Vimentin and p-S6 and the percentage of p-S6+ DCCs in clusters (more than 3 and less than 9) was plotted (N= 116 DCCs in shC, N= 103 DCCs in shC C26).. Mann Whitney.

| Follow-up | Total Patients | Patients with Dominant DCC phenotypes in LN | | | |
|---|---|---|---|---|---|
| | | Reactivating MDK$^{HIGH}$/ NR2F1$^{LOW}$ | Dormant MDK$^{LOW}$/ NR2F1$^{HIGH}$ | Double Negative | MDK$^{LOW}$/ NR2F1$^{HIGH}$ Double Negative (1:1) |
| Deceased (<3 years) | 4 | 3 (75%) | 0 (0%) | 0 (0%) | 1 (25%) |
| Alive/Recurred (>7 years) | 6 | 0 (0%) | 1 (16%) | 2 (33%) | 3 (50%) |

**Supplementary Table 1.** Small cohort of patients with follow up information shows that 3 out of 4 patients that were deceased in less than 3 years carried a dominant putative reactivating MDK$^{HIGH}$/NR2F1$^{LOW}$ DCC population (75% of patients). On the other hand, 6 out of 6 patients that remained alive 6-19 years later or recurred after 7 years carried a dominant putative dormant MDK$^{LOW}$/NR2F1$^{HIGH}$ DCC, dominant double negative, or half MDK$^{LOW}$/NR2F1$^{HIGH}$ and half double negative phenotypes.
